# PRMT7 mediated PTEN activation promotes bone formation in female mice

**DOI:** 10.1101/2024.07.31.605998

**Authors:** Yingfei Zhang, Jia Qing, Yang Li, Xin Gao, Dazhuang Lu, Yiyang Wang, Lanxin Gu, Hui Zhang, Zechuan Li, Xu Wang, Yongsheng Zhou, Ping Zhang

## Abstract

Although the epigenetic mechanisms underlying bone formation are recognized, their specific roles and regulatory pathways remain largely unexplored. In this study, we unveil PRMT7 as a novel epigenetic modulator of MSCs’ osteogenic commitment. The conditional knockout of *Prmt7* in mice reveals significantly impaired osteogenesis and bone regeneration exclusively in females, affecting both long bones and craniofacial structures, with no discernible impact in males. Our findings demonstrate that PRMT7 orchestrates osteogenic differentiation through a methyltransferase-dependent manner. Mechanistically, PRMT7 modulates MSCs’ osteogenic differentiation through the activation of PTEN. Specifically, PRMT7 augments *PTEN* transcription by increasing H3R2me1 levels at the *PTEN* promoter. Furthermore, PRMT7 interacts with the PTEN protein, and its deficiency leads to the ubiquitination and subsequent degradation of nuclear PTEN, revealing an unprecedented pathway. Crucially, PTEN overexpression ameliorates the osteogenic deficits observed in *Prmt7*-deficient mice. Our research positions PRMT7 as a potential therapeutic target to enhance bone formation and offers novel molecular insights into the PRMT7-PTEN regulatory axis, underscoring its significance in bone biology and regenerative medicine.

**Subject Categories** Developmental Biology, Musculoskeletal System, Epigenetics, Post-translational Modifications

## Introduction

Epigenetic modifications, particularly histone modifications, play a crucial role in regulating cell pluripotency by influencing chromatin condensation and the transcription of bone-related genes (*Xu et al., 2020; Montecino et al., 2020*). Studies predominantly indicate that H3K4me3, a marker of gene activation, actively promotes osteogenic differentiation, whereas H3K9me3 and H3K27me3 contribute to more condensed heterochromatin formation, thereby inhibiting the expression of osteogenic genes (*Rojas et al., 2019; Nicetto et al., 2019; Ye et al., 2012; Qi et al., 2020; Jing et al., 2016; Wei et al., 2011*). However, the role of histone arginine methylation in osteogenesis remains under-investigated.

Histone and non-histone arginine methylation, mediated by protein arginine methyltransferases (PRMTs), are essential for various biological functions, including DNA repair, gene transcription, signal transduction, and stem cell fate determination (*Guccione & Richard, 2019*). PRMTs, which methylate arginine residues on proteins, are categorized into three types based on their methylated arginine products. Type I PRMTs (PRMT-1, 2, 3, 4, 6, and 8) catalyze asymmetric dimethylarginine (ADMA); Type II PRMTs (PRMT-5, 9) catalyze symmetric dimethylarginine (SDMA); and PRMT7, the sole Type III methyltransferase, functions exclusively as a monomethylarginine (MMA) transferase (*Tewary et al., 2019*). Our research, along with others, has demonstrated the significant role of PRMTs in MSC-dependent osteogenesis. For instance, PRMT3 enhances miR-3648 expression by increasing H4R3me2a levels, thereby accelerating MSC osteoblastic differentiation (*Min et al., 2019*). CARM1/PRMT4 facilitates MSC osteogenic differentiation by enhancing various H3 methylation sites (*Li et al., 2023a; Jo et al., 2012*). Inhibition of PRMT5 results in a global downregulation of H4R3me2s and H3R8me2s, particularly at the promoters of *Gbp2* and *Gbp5*, promoting MSC osteogenic differentiation (*Kota et al., 2018*). PRMT6 positively regulates MSC osteogenic differentiation, although specific histone modifications were not detailed (*Li et al., 2021b*). Given these findings, we hypothesize that PRMT7 can also significantly influence MSC-mediated osteogenesis.

Histone post-translational modifications (PTMs), as a “histone code” mediated by the PTM writer (PRMTs are encompassed within this category), reader, and eraser, fall within the realm of epigenetics, regulating genomic response (*Strahl & Allis, 2000; Millán-Zambrano et al., 2022*). Meanwhile, the PTMs of non-histone substrates, akin to histone PTMs, also play a pivotal role in various biological processes through a similar “writer-reader-eraser” axis (*Narita et al., 2019; Biggar & Li, 2015*). PTEN, a tumor suppressor, regulates various biological functions through its intricate histone and PTEN codes, influencing transcription and PTMs of itself individually, subsequently leading to alterations in PTEN expression level (*Lee et al., 2018; González-García et al., 2022; Chen et al., 2020*). Current research reveals that changes in histone acetylation and methylation at the *PTEN* promoter can impact its expression (*Shen et al., 2019; Lu et al., 2009*). Moreover, PTMs such as phosphorylation (*Adey et al., 2000; Rahdar et al., 2009; Fenton et al., 2012; Vazquez et al., 2000*), acetylation (*Okumura et al., 2006; Ikenoue et al., 2008; Meng et al., 2016*), methylation (*Feng et al., 2019*), and ubiquitination significantly contribute to PTEN loss in various cancers. In particular, PTEN degradation via the ubiquitin-mediated proteasome system is crucial for maintaining its expression level (*Wang et al., 2007; Trotman et al., 2007a; Fouladkou et al., 2008; Lee et al., 2019*). Variations in PTEN expression can precipitate a spectrum of hereditary disorders, cancers, and other diseases (*Lee et al., 2018*).

Here, we demonstrate that PRMT7 exclusively promotes osteogenesis and bone regeneration in both axial and appendicular bones in female mice. PRMT7 regulates MSC osteogenic differentiation by activating the target gene *PTEN* through dual mechanisms: 1) PRMT7 binds to the *PTEN* promoter, enhancing H3R2me1 levels to transcriptionally activate PTEN; 2) PRMT7 interacts with the PTEN protein, stabilizing nuclear PTEN through a non-methyltransferase-dependent pathway. Osteopenia resulting from PRMT7 deletion can be rescued by AAV-delivered *Pten*. In summary, our study demonstrates that PRMT7 positively regulates osteogenesis by transcriptionally activating PTEN and stabilizing nuclear PTEN, offering a novel therapeutic target for bone regenerative medicine.

## Results

### Deletion of *Prmt7* impairs long bone development and regeneration

To investigate the role of PRMT7 in bone formation *in vivo*, we employed two distinct Cre mice to establish *Prmt7* conditional knockout (CKO) mouse models. These models were developed by crossing *Prmt7^f/f^* mice with *Prrx1-Cre* mice (*Prrx1-Cre; Prmt7^f/f^*) and *Sp7-Cre* mice (*Sp7-Cre; Prmt7^f/f^*) individually (Appendix Fig. 1A-B). At 6 weeks of age, female *Prrx1-Cre; Prmt7^f/f^* and *Sp7-Cre; Prmt7^f/f^* mice exhibited dwarfism compared to their control littermates (*Prmt7^f/f^*), which was not observed in male mice (Fig. 1A). Alizarin red and Alcian blue staining displayed remarkably delayed bone formation in *Prrx1-Cre; Prmt7^f/f^*and *Sp7-Cre; Prmt7^f/f^* female mice at E18.5d and P0d compared to control *Prmt7^f/f^* littermates (Fig. 1B). This delay was not observed in males (Appendix Fig. 1C). Both *Prmt7* CKO female mice exhibited decreased bone mineral density (BMD), bone volume (BV/TV), trabecular number (Tb. N), and increased trabecular space (Tb. Sp) (Fig. 1C-E), with no significant differences in males (Appendix Fig. 1D-F).

**Figure 1.**
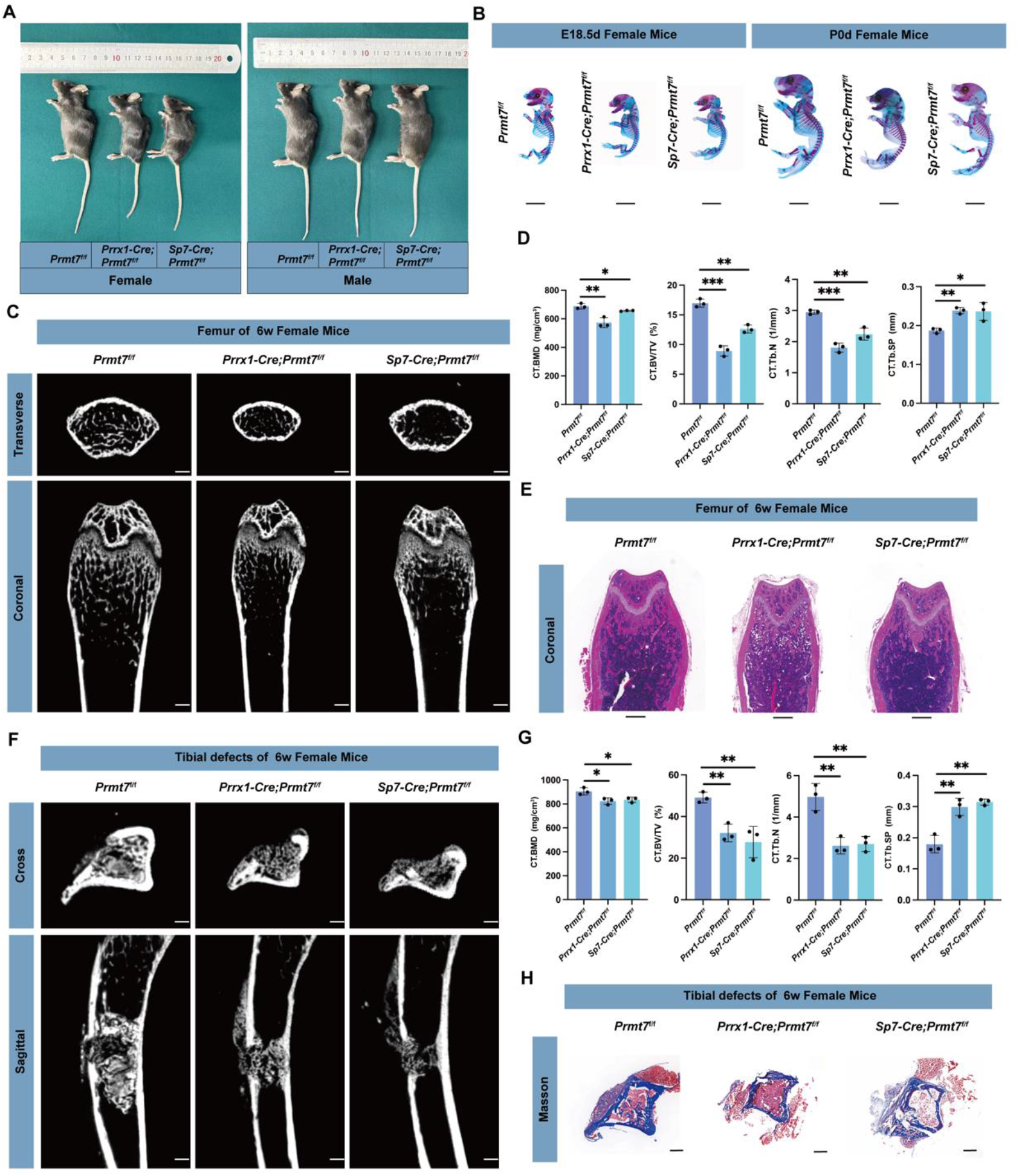
PRMT7 deficiency affects long bone development and regeneration in female mice. A. Overall physical size of 6-week-old female and male control and CKO mice (*Prrx1-cre; Prmt7^f/f^*and *Sp7-cre; Prmt7^f/f^*). B. Alcian Blue and Alizarin Red staining of E18.5 and P0 female and male control and CKO mice (*Prrx1-cre; Prmt7^f/f^* and *Sp7-cre; Prmt7^f/f^*). Scale bar: 5 mm. C. Micro-CT analysis of femurs from 6-week-old female control and CKO mice (*Prrx1-cre; Prmt7^f/f^* and *Sp7-cre; Prmt7^f/f^*). The upper panel shows the cross-sectional view of the metaphysis; the lower panel shows the coronal view of the metaphysis. Scale bar: 500 μm. D. Quantitative analysis of bone parameters at the femoral metaphysis growth plate, including BMD, BV/TV, Tb. N, and Tb. Sp, obtained in C (n = 3 mice for each genotype). E. H&E staining of the coronal section of the femur in 6-week-old female control and CKO mice (*Prrx1-cre; Prmt7^f/f^* and *Sp7-cre; Prmt7^f/f^*). Scale bar: 500 μm. F. Micro-CT analysis of tibial defect modeling in 6-week-old female control and CKO mice (*Prrx1-cre; Prmt7^f/f^* and *Sp7-cre; Prmt7^f/f^*), with samples collected 10 days post-operation. The upper panel shows the cross-sectional view of the defect site; the lower panel shows the sagittal view of the tibia sample. Scale bar: 500 μm. G. Quantitative analysis of bone parameters in the defect region of the tibial defect model obtained in F (n = 3 mice for each genotype). H. Cross-sectional Masson’s trichrome staining of the obtained tibial defect models in 6-week-old female mice. Scale bar: 500 μm. In D and G, data are presented as mean ± SD (*, *P*<0.05, **, *P* <0.01, ***, *P* <0.001) (One-way ANOVA).

To explore the bone regeneration capacity of PRMT7, we established a tibial defect model in 6-week-old *Prmt7* CKO mice, collecting samples 10 days post-injury. Micro-CT analysis showed poor cortical bone healing, larger bone gaps, and reduced trabecular bone formation in both *Prmt7* CKO strains compared to control *Prmt7^f/f^*littermates (Fig. 1F). Quantitative analysis of bone parameters around the defect area demonstrated a significant decrease in BMD, BV/TV, and Tb. N, and an increase in Tb. Sp in *Prmt7* CKO female mice, changes that were absent in males (Fig. 1G, Appendix Fig. 1G-H). Masson staining corroborated these findings, showing less new bone formation in the knockouts compared to the control group (Fig. 1H). These phenotypes were not observed in male mice (Appendix Fig. 1I). In conclusion, *Prmt7* deficiency impairs long bone formation and bone regeneration ability in female mice.

### *Prmt7* CKO mice exhibit impaired craniofacial bone and dental structures

Unlike appendicular bones, craniofacial bones have distinct developmental origins, osteogenesis patterns, and exist in a more complex anatomical and physiological environment. Teeth, similar to bones, function as biomineralizing tissues attached to the craniomaxillofacial structure (Salhotra et al., 2020; Berendsen & Olsen, 2015). We investigated whether PRMT7 affects the structure of craniofacial bone and dental structures. Micro-CT and H&E analyses revealed that *Prmt7*-deficient mice had smaller skull volumes, thinner bones, and less homogeneous bone structure. Quantitative analysis of bone parameters showed reduced BMD, BV/TV, and Tb. N in *Prmt7*-deficient female mice, along with increased trabecular space (Tb. Sp) (Fig. 2A-C). Consistently, these phenomena were observed exclusively in female mice and not in male mice (Appendix Fig. 2A-C). Furthermore, Micro-CT of the mandibles of both *Prmt7*-deficient strains indicated reduced periapical bone mass around the mandibular first molar and a thinner bone wall of the mandibular body in female mice, accompanied by the same trend in bone parameters as the morphology (Fig. 2D-E). H&E staining showed significantly reduced cementum at the apex of the mandibular first molar, and toluidine blue staining showed widened pre-dentin in the distal mandibular first molar, indicating poor dentin mineralization (Fig. 2F). These phenotypes were not observed in male mice (Appendix Fig. 2D-F).

**Figure 2.**
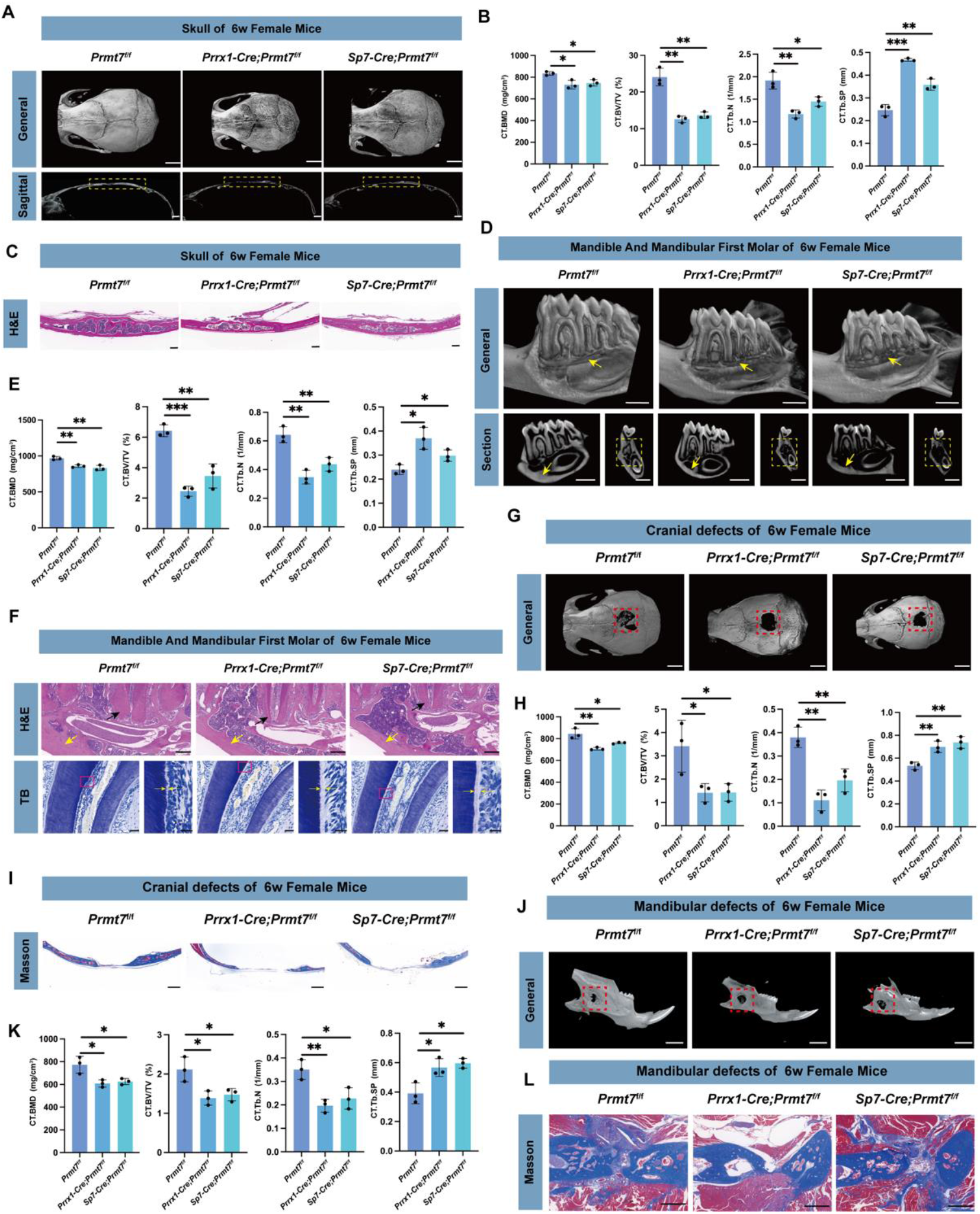
PRMT7 deficiency impairs craniofacial bone and dental structures in female mice. A. Micro-CT analysis of the skull in 6-week-old female control and CKO mice (*Prrx1-cre; Prmt7^f/f^* and *Sp7-cre; Prmt7^f/f^*). The upper panel: 3D reconstruction of the whole skull; The lower panel: midsagittal view of the skull. The yellow dashed box indicates the calvarial region. Scale bars:2 mm (upper) and 1 mm (lower). B. Quantitative analysis of bone parameters at the calvarial region in 6-week-old female control and CKO mice (*Prrx1-cre; Prmt7^f/f^* and *Sp7-cre; Prmt7^f/f^*), including BMD, BV/TV, Tb. N, and Tb. Sp, obtained in A (n = 3 mice for each genotype). C. H&E staining of the midcoronal sections of the skull in 6-week-old female control and CKO mice (*Prrx1-cre; Prmt7^f/f^* and *Sp7-cre; Prmt7^f/f^*). Scale bar: 200 μm. D. Micro-CT analysis of the mandible in 6-week-old female control and CKO mice (*Prrx1-cre; Prmt7^f/f^* and *Sp7-cre; Prmt7^f/f^*). The upper panel: 3D reconstructed sectional view of the mandible; The lower left panel: mesiodistal sectional view of the first molar; The lower right panel: coronal sectional view of the first molar. The yellow arrow at the upper panel points to the bone around the apical area of the distal root of the first molar; the yellow arrow at the lower left panel points to the bone wall at the lower border of the mandibular body; the yellow dashed box at the lower right panel indicates the bone between the furcation of the first molar and the mandibular canal. Scale bars: 1 mm. E. Bone parameters quantitative analysis of the mandible in 6-week-old female control and CKO mice (*Prrx1-cre; Prmt7^f/f^* and *Sp7-cre; Prmt7^f/f^*), including BMD, BV/TV, Tb. N, and Tb. Sp, obtained in A (n = 3 mice for each genotype). F. H&E staining (The upper panel) of the periapical bone in the mandible of 6-week-old female control and CKO mice (*Prrx1-cre; Prmt7^f/f^* and *Sp7-cre; Prmt7^f/f^*). The black arrow indicates the cementum in the apical area of the distal root of the first molar, while the yellow arrow points to the bone wall at the lower border of the mandible. Toluidine Blue staining (the lower panel) of the dental tissues at the alveolar crest of the distal root of the first molar. The yellow arrow in the lower right points to the predentin, which is an enlarged view of the area within the pink box in the lower left. Scale bars:200 μm (upper), 50 μm (lower left) and 20 μm (lower right). G. Micro-CT analysis of the cranial defect model in 6-week-old female control and CKO mice (*Prrx1-cre; Prmt7^f/f^*and *Sp7-cre; Prmt7^f/f^*). The red dashed box indicates the defect area. scale bar: 2 mm. H. Quantitative analysis of bone parameters in the defect region of the cranial defect model obtained in G (n = 3 mice for each genotype). I. Coronal-sectional Masson’s trichrome staining of the obtained cranial defect models in 6-week-old female control and CKO mice (*Prrx1-cre; Prmt7^f/f^* and *Sp7-cre; Prmt7^f/f^*). Scale bar: 500 μm. J. Micro-CT analysis of the mandibular defect model in 6-week-old female control and CKO mice (*Prrx1-cre; Prmt7^f/f^* and *Sp7-cre; Prmt7^f/f^*). The red dashed box indicates the defect area. scale bar: 2 mm. K. Quantitative analysis of bone parameters in the defect region of the mandibular defect model obtained in J (n = 3 mice for each genotype). L. Cross-sectional Masson’s trichrome staining of the obtained mandibular defect models in 6-week-old female control and CKO mice (*Prrx1-cre; Prmt7^f/f^* and *Sp7-cre; Prmt7^f/f^*). Scale bar: 500 μm. In B, E, H and K data are presented as mean ± SD (*, *P* <0.05, **, *P* <0.01, ***, *P* <0.001) (One-way ANOVA).

To confirm whether PRMT7 deficiency impairs craniofacial bone regeneration, we established skull and mandible injury models and harvested samples after two months. In control mice, the 2-mm diameter defect was nearly healed, whereas in CKO female mice, the defect remained large. Quantification of bone parameters demonstrated lower BMD, reduced BV/TV, and less bone mass in the defect area in *Prmt7* CKO female mice (Fig. 2G-I). Male mice did not exhibit changes in calvarial bone regeneration capacity (Appendix Fig. 2G-I). A consistent phenotype was observed in the mandibular body defect model in female mice. Micro-CT and Masson staining showed that the defects in control mice were nearly healed, whereas those in CKO mice remained significantly larger. Quantification analysis confirmed these phenotypic differences (Fig. 2J-L). Mandibular regeneration capacity was not affected in males (Appendix Fig. 2J-L). The above results show that PRMT7 deficiency impairs craniofacial bone and dental structures and weakens craniofacial bone regeneration.

### PRMT7 regulates osteogenic differentiation in a methyltransferase activity-dependent manner

To explore the effect of PRMT7 on the osteogenic differentiation of BMSCs, we extracted BMSCs from female *Prmt7^f/f^*, *Prrx1-Cre; Prmt7^f/f^* and *Sp7-Cre; Prmt7^f/f^* mice (mBMSCs) for subsequent functional experiments. Western blot and qRT-PCR were used to validate the knockout efficiency of PRMT7 (Fig. 3A; Appendix Fig. 3A). ALP activity analysis revealed that the osteogenic differentiation ability of mBMSCs from *Prmt7*-deficient mice was significantly decreased compared to the control group (Fig. 3B). Additionally, western blot and qRT-PCR demonstrated that the expression of RUNX2, a transcription factor associated with osteogenic differentiation, was significantly decreased in *Prmt7*-deficient mBMSCs following osteogenic induction (Fig. 3C-D; Appendix Fig. 3B). Next, human BMSCs (hBMSCs) were induced for osteogenesis over 7 days and 14 days. The levels of PRMT7 protein and mRNA increased with the duration of osteogenic induction (Appendix Fig. 3C-D). To further confirm the critical role of PRMT7 in human cells, we established PRMT7 stable knockdown cells. The efficiency of PRMT7 knockdown was confirmed by western blot and qRT-PCR analysis (Fig. 3E; Appendix Fig. 3E). After 7 days of osteogenic induction in both control and PRMT7 knockdown cells, ALP staining and quantitative analysis showed that the osteogenic differentiation ability was reduced in PRMT7 knockdown cells (Fig. 3F). Figure 3G-H and Appendix Fig. 3F displayed that PRMT7 knockdown hBMSCs had significantly lower levels of RUNX2 after induction compared with the control group, indicating decreased osteogenic differentiation ability. In summary, PRMT7 promotes the osteogenic differentiation of BMSCs *in vitro*.

**Figure 3.**
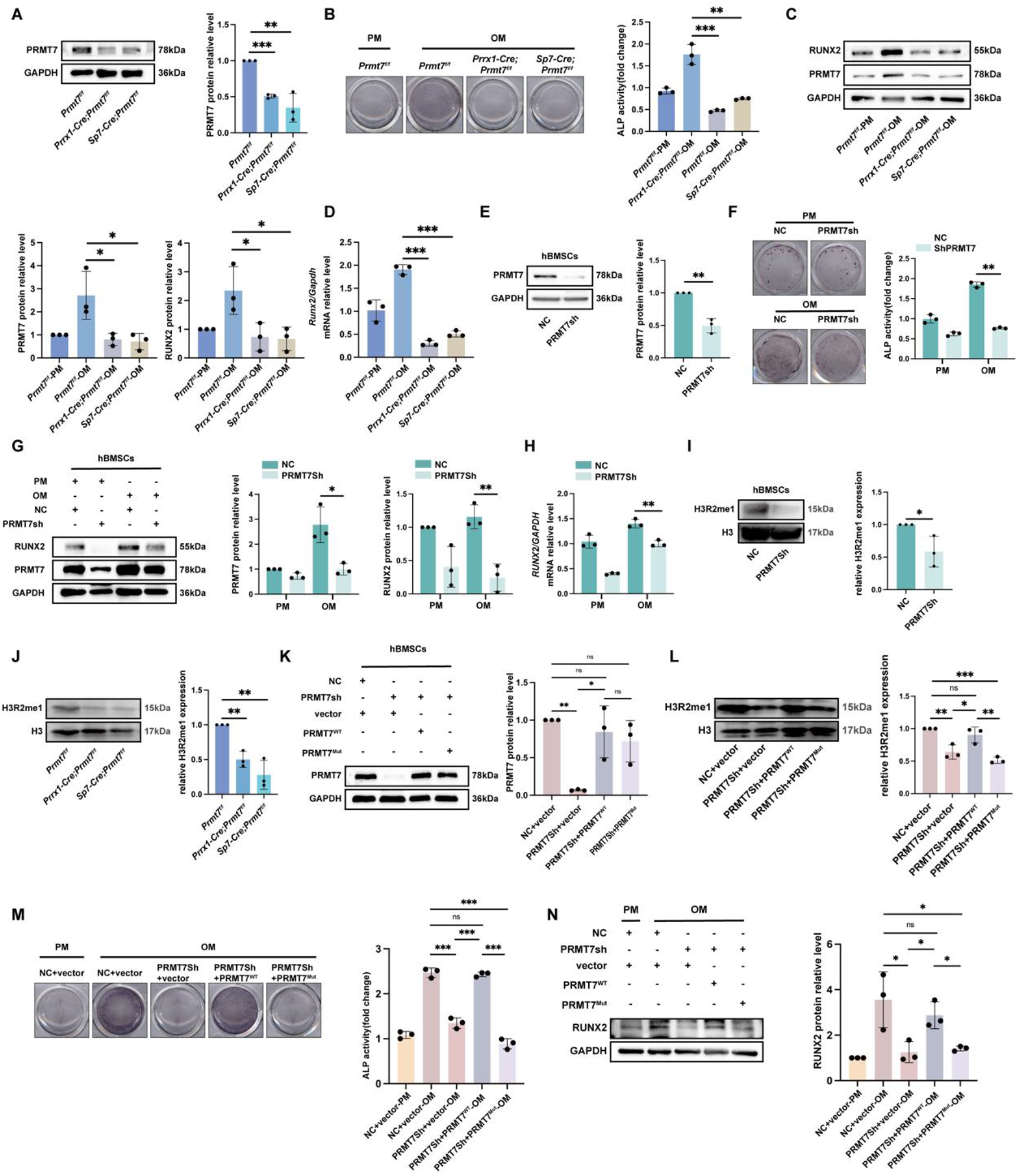
PRMT7 regulates osteogenic differentiation via its methyltransferase activity. A. Representative Western blot images (left) of PRMT7 in female *Prmt7^f/f^*, *Prrx1-Cre; Prmt7^f/f^* and *Sp7-Cre; Prmt7^f/f^* mBMSCs. Quantification (right) of relative PRMT7 levels normalized to GAPDH. B. Staining and quantification of ALP in female *Prmt7^f/f^*, *Prrx1-Cre; Prmt7^f/f^* and *Sp7-Cre; Prmt7^f/f^* mBMSCs after 7 days of osteogenic induction. C. Representative Western blot images (left) of PRMT7 and RUNX2 in female *Prmt7^f/f^*, *Prrx1-Cre; Prmt7^f/f^* and *Sp7-Cre; Prmt7^f/f^* mBMSCs after 7 days of osteogenic induction. Quantification (right) of relative PRMT7 and RUNX2 levels normalized to GAPDH. D. Representative qRT-PCR results of *Runx2* in female *Prmt7^f/f^*, *Prrx1-Cre; Prmt7^f/f^* and *Sp7-Cre; Prmt7^f/f^* mBMSCs after 7 days of osteogenic induction. E. Representative Western blot images (left) of PRMT7 in hBMSCs after transfection with PRMT7sh and control lentivirus respectively. Quantification (right) of relative PRMT7 levels normalized to GAPDH. F. Staining and quantification of ALP in PRMT7sh and control hBMSCs after 7 days of osteogenic induction. G. Representative Western blot images (left) of RUNX2 and PRMT7 in PRMT7sh and control hBMSCs after 7 days of osteogenic induction. Quantification (right) of relative RUNX2 and PRMT7 levels normalized to GAPDH. H. Representative qRT-PCR results of *RUNX2* in PRMT7sh and control hBMSCs after 7 days of osteogenic induction. I. Representative Western blot images (upper) of H3R2me1 in PRMT7sh and control hBMSCs after extracting nuclear proteins. Quantification (lower) of relative H3R2me1 levels normalized to H3. J. Representative Western blot images (The upper) of H3R2me1 in female *Prmt7^f/f^*, *Prrx1-Cre; Prmt7^f/f^* and *Sp7-Cre; Prmt7^f/f^* mBMSCs after extracting nuclear proteins. Quantification (the lower) of relative H3R2me1 levels normalized to H3. K. Representative Western blot images (The upper) of PRMT7 in hBMSCs after transfection with lentivirus carrying either an empty vector, wildtype PRMT7 (PRMT7^WT^), or enzyme-mutant PRMT7 (PRMT7^Mut^). Quantification (the lower) of relative PRMT7 levels normalized to GAPDH. L. Representative Western blot images (left) of H3R2me1 after transfection with lentivirus carrying either an empty vector, wildtype PRMT7 (PRMT7^WT^), or enzyme-mutant PRMT7 (PRMT7^Mut^) in hBMSCs and extraction of nuclear proteins. Quantification (right) of relative H3R2me1 levels normalized to H3. M. Staining and quantification of ALP in hBMSCs after transfection with lentivirus carrying either an empty vector, wildtype PRMT7 (PRMT7^WT^), or enzyme-mutant PRMT7 (PRMT7^Mut^) and 7 days of osteogenic induction. N. Representative Western blot images (left) of RUNX2 in hBMSCs after transfection with lentivirus carrying either an empty vector, wildtype PRMT7 (PRMT7^WT^), or enzyme-mutant PRMT7 (PRMT7^Mut^) and 7 days of osteogenic induction. Quantification (right) of relative RUNX2 levels normalized to GAPDH. All data are mean ± SD, n = 3 biological replicates. (ns, not significant, *, *P* <0.05, **, *P* <0.01, ***, *P* <0.001) (Independent samples t-test or One-way ANOVA).

To probe whether PRMT7 regulates osteogenic differentiation in an enzyme activity-dependent manner, we utilized a plasmid with already reported mutations in the PRMT7 enzyme active site (E144A, D147A, E153A) *(Li et al., 2021a*). PRMT7 is known to modulate H3R2me1 levels in MH-S cells (*Günes Günsel et al., 2022*). Thus, we initially investigated whether the trends in H3R2me1 levels in hBMSCs and mBMSCs align with those observed in MH-S cells. Western blot analysis revealed that silencing PRMT7 resulted in a significant decrease in the global levels of H3R2me1 in MSCs (Fig. 3I-J). Then, we transfected hBMSCs with lentivirus carrying either an empty vector, wildtype PRMT7 (PRMT7^WT^), or enzyme-mutant PRMT7 (PRMT7^Mut^). Western blot confirmed the re-introduction of PRMT7 (Fig. 3K). Figure L demonstrates that the decrease in H3R2me1 levels induced by PRMT7 knockdown cannot be rescued by PRMT7^Mut^, thereby confirming the validity of the enzymatic activity mutation. Osteogenic induction was performed on different groups of hBMSCs. We found that the addition of WT-PRMT7 plasmid to the PRMT7 knockdown cells restored osteogenic differentiation ability. In contrast, the PRMT7^Mut^ plasmid did not improve the osteogenic differentiation ability of PRMT7 knockdown cells (Fig. 3M-N). This leads to the conclusion that PRMT7 promotes the osteogenic differentiation of BMSCs in an enzyme activity-dependent manner.

### PRMT7 activates PTEN in female mice

To reveal the mechanism by which PRMT7 regulates osteogenic differentiation, we performed RNA sequencing on mBMSCs from CKO mice of both strains and compared them with the control mice. We identified 354 down-regulated genes and 554 up-regulated genes in *Prrx1-Cre; Prmt7^f/f^* and *Prmt7^f/f^* mice. Gene Ontology (GO) analysis showed that the enriched functions were primarily related to regeneration and tooth mineralization, while the KEGG pathway enrichment highlighted the PI3K-AKT signaling pathway (Appendix Fig. 4A-C). In the comparison between *Sp7-Cre; Prmt7^f/f^*and *Prmt7^f/f^* mice, we found 658 differentially expressed genes (DEGs). GO pathway enrichment was mainly in the Wnt signaling pathway and mesenchymal cell differentiation, and KEGG pathway enrichment was the mTOR signaling pathway (Appendix Fig. 4D-F). We then merged the two groups of DEGs, identifying a total of 10 DEGs, which were visualized in heat maps (Fig. 4A-C). We screened for genes with consistent changes in both heat maps and validated them by RT-qPCR. Only *Pten* showed a consistent trend with the sequencing data, being down-regulated in both lines compared to the control group (Fig. 4D). Next, we further examined whether PTEN expression was consistent in different cell lines. In female mBMSCs, PTEN protein level consistent with mRNA tendency was significantly reduced in CKO mice (Fig. 4E). However, in male mice mBMSCs, PTEN expression remained unchanged at both the protein and mRNA levels (Fig. 4F-G). These findings suggest that PTEN may contribute to the gender differences observed in the phenotypes of *Prmt7*-deficient mice. Meanwhile, we also detected the PTEN expression in hBMSCs and dental pulp stem cells (DPSCs); protein and mRNA levels showed that knockdown of PRMT7 caused PTEN expression level reduction (Fig. 4H-I; Appendix Fig. 4G-H), while overexpression of PRMT7 showed the opposite trend (Appendix Fig. 4I-L). Next, our results implicated that PRMT7 activated PTEN in an enzyme-dependent manner, as demonstrated using PRMT7^WT^ and PRMT7^Mut^ plasmids in hBMSCs (Fig. 4J). In summary, we demonstrated that PRMT7 activates PTEN in female mice through an enzymatic activity-dependent mechanism.

**Figure 4.**
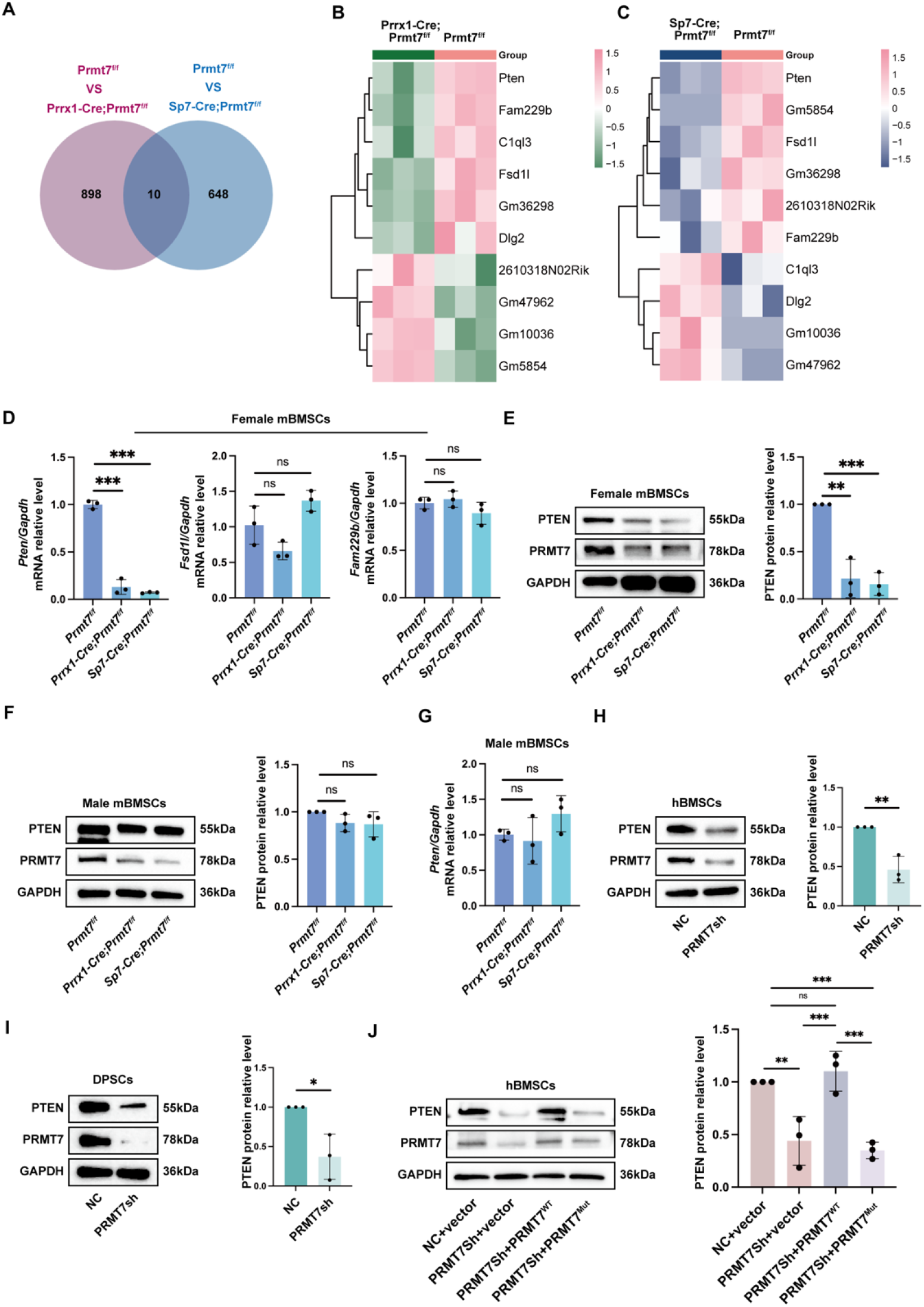
PRMT7 induces PTEN activation in female mice. A. Venn diagram illustrating the overlap of differentially expressed genes between control and two strains CKO groups respectively, highlighting shared and unique genes in each group. B. Heatmap displaying the expression levels of DEGs between *Prmt7^f/f^* and *Prrx1-Cre; Prmt7^f/f^* groups, with color intensity representing the level of gene expression across samples. C. Heatmap displaying the expression levels of DEGs between *Prmt7^f/f^* and *Sp7-Cre; Prmt7^f/f^* groups, with color intensity representing the level of gene expression across samples. D. Representative qRT-PCR results of *Pten*, *Fsd1l*, *Fam229b* in female *Prmt7^f/f^*, *Prrx1-Cre; Prmt7^f/f^* and *Sp7-Cre; Prmt7^f/f^* mBMSCs. E. Representative Western blot images (left) of PTEN in female *Prmt7^f/f^*, *Prrx1-Cre; Prmt7^f/f^*and *Sp7-Cre; Prmt7^f/f^* mBMSCs. Quantification (right) of relative PTEN levels normalized to GAPDH. F. Representative Western blot images (left) of PTEN in male *Prmt7^f/f^*, *Prrx1-Cre; Prmt7^f/f^*and *Sp7-Cre; Prmt7^f/f^* mBMSCs. Quantification (right) of relative PTEN levels normalized to GAPDH. G. Representative qRT-PCR results of *Pten* in male *Prmt7^f/f^*, *Prrx1-Cre; Prmt7^f/f^* and *Sp7-Cre; Prmt7^f/f^* mBMSCs. H. Representative Western blot images (left) of PTEN in PRMT7sh and control hBMSCs. Quantification (right) of relative PTEN levels normalized to GAPDH. I. Representative Western blot images (left) of PTEN in PRMT7sh and control DPSCs. Quantification (right) of relative PTEN levels normalized to GAPDH. J. Representative Western blot images (left) of PTEN in hBMSCs after transfection with lentivirus carrying either an empty vector, wildtype PRMT7 (PRMT7^WT^), or enzyme-mutant PRMT7 (PRMT7^Mut^) and 7 days of osteogenic induction. Quantification (right) of relative PTEN levels normalized to GAPDH. All data are mean ± SD, n = 3 biological replicates. (ns, not significant, *, *P* <0.05, **, *P* <0.01, ***, *P* <0.001) (Independent samples t-test or One-way ANOVA).

### PRMT7 acts as an H3R2me1 histone methyltransferase in *PTEN* promoter

As a protein arginine methyltransferase, PRMT7 modifies histone methylation and influences the transcriptional regulation of target genes (*Günes Günsel et al., 2022*). To probe the regulatory role of PRMT7 in PTEN expression through histone methylation at the *PTEN* promoter region, we first conducted ChIP-qPCR. We designed primers for the *PTEN* promoter region upstream of the *PTEN* transcription start site (TSS) according to a putative enhancer (Fig. 5A). Our results indicated that PRMT7 binds to the promoter region of *PTEN*, while PRMT7 occupancy on the promoters of *PTEN* was reduced in PRMT7 knockdown cells, confirming PRMT7’s binding (Fig. 5B). Meanwhile, the same results were validated in female *Prmt7^f/f^*, *Prrx1-Cre; Prmt7^f/f^* and *Sp7-Cre; Prmt7^f/f^* mBMSCs (Fig. 5C). Additionally, decreased binding of PRMT7 at the promoter regions was associated with decreased occurrence of its substrate, H3R2me1, which is a hallmark of gene activation (Fig. 5D) (Kirmizis et al., 2009). In *Prmt7*-knockout mice compared to controls, there was also a reduction in H3R2me1 levels at the *Pten* promoter region (Fig. 5E). These results indicate that PRMT7 transcriptionally activates *PTEN* via monomethylation of H3R2. To determine whether PRMT7 regulates *PTEN* transcriptional activation in an enzyme activity-dependent manner, we introduced a PRMT7 enzyme inactivation mutant plasmid into PRMT7 knockdown cells. ChIP-qPCR results showed that this mutant did not restore H3R2me1 levels in the *PTEN* promoter region (Fig. 5F). The above results show that PRMT7, functioning as an H3R2 monomethyl transferase, mediates *PTEN* transcription activation in an enzyme activity-dependent manner.

**Figure 5:**
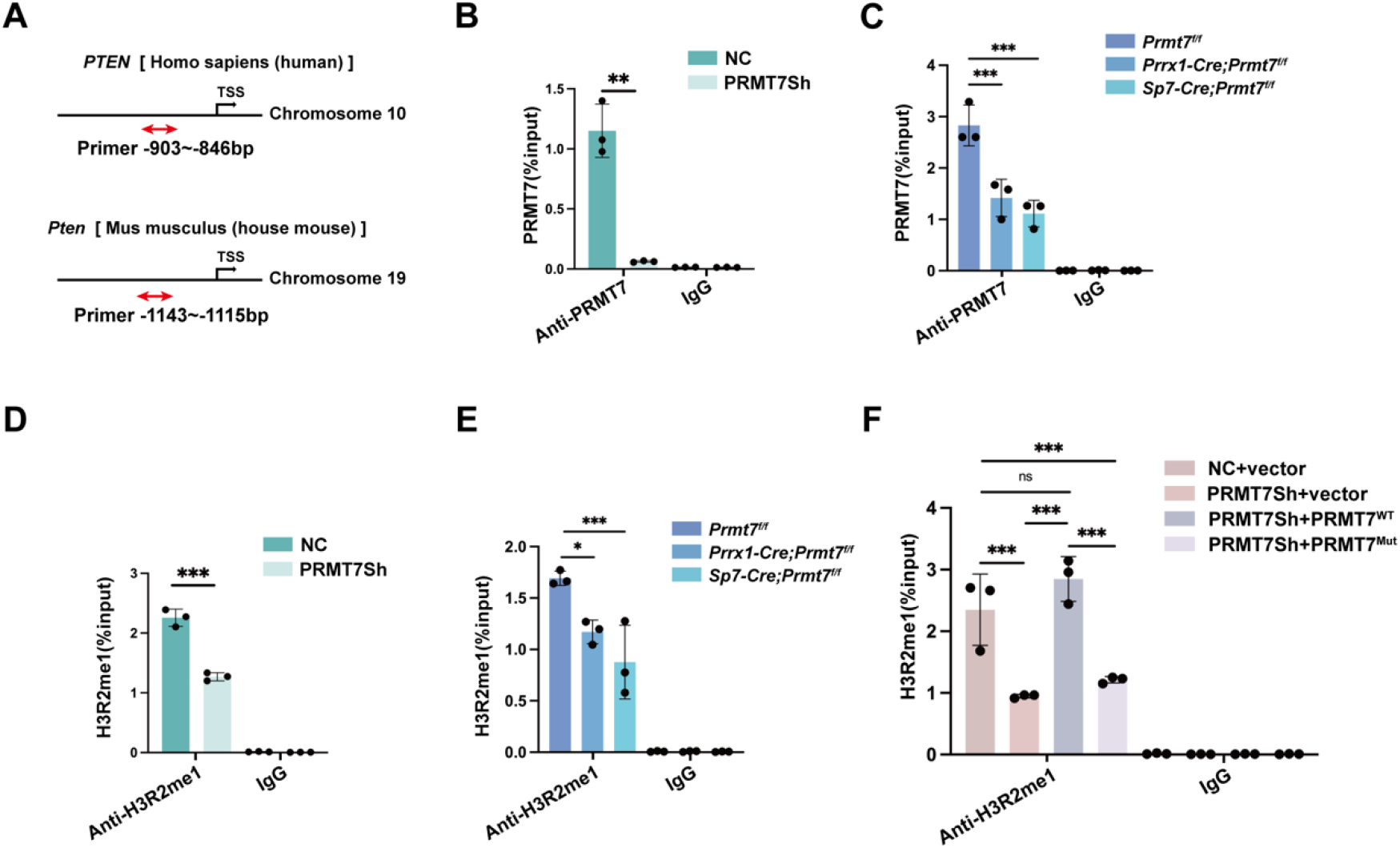
PRMT7 specifically methylates H3R2me1 at the *PTEN* promoter. A. Structural schematic of human *PTEN* and mouse *Pten* genes. Red bidirectional arrows indicate regions detected by ChIP. TSS: Transcription Start Site. B. The ChIP experiments were performed using anti-PRMT7 antibodies on PRMT7sh and control hBMSCs. C. The ChIP experiments were performed using anti-PRMT7 antibodies on female *Prmt7^f/f^*, *Prrx1-Cre; Prmt7^f/f^* and *Sp7-Cre; Prmt7^f/f^* mBMSCs. D. The ChIP experiments were performed using anti-H3R2me1 antibodies on PRMT7sh and control hBMSCs. E. The ChIP experiments were performed using anti-H3R2me1 antibodies on female *Prmt7^f/f^*, *Prrx1-Cre; Prmt7^f/f^* and *Sp7-Cre; Prmt7^f/f^*mBMSCs. F. The ChIP experiments were performed using anti-H3R2me1 antibodies on hBMSCs after transfection with lentivirus carrying either an empty vector, wildtype PRMT7 (PRMT7^WT^), or enzyme-mutant PRMT7 (PRMT7^Mut^). All data are mean ± SD, n = 3 biological replicates. (ns, not significant, *, *P* <0.05, **, *P* <0.01, ***, *P* <0.001) (One-way ANOVA).

### PRMT7 interacts with and stabilizes nuclear PTEN

To further explore the relationship between PRMT7 and PTEN, we performed forward and reverse Co-IP to confirm the interaction between PRMT7 and PTEN in HEK293T and hBMSCs (Fig. 6A). Additionally, we used immunofluorescence to confirm the colocalization of PRMT7 and PTEN in different cell types (Fig. 6B; Appendix Fig. 5A).

**Figure 6.**
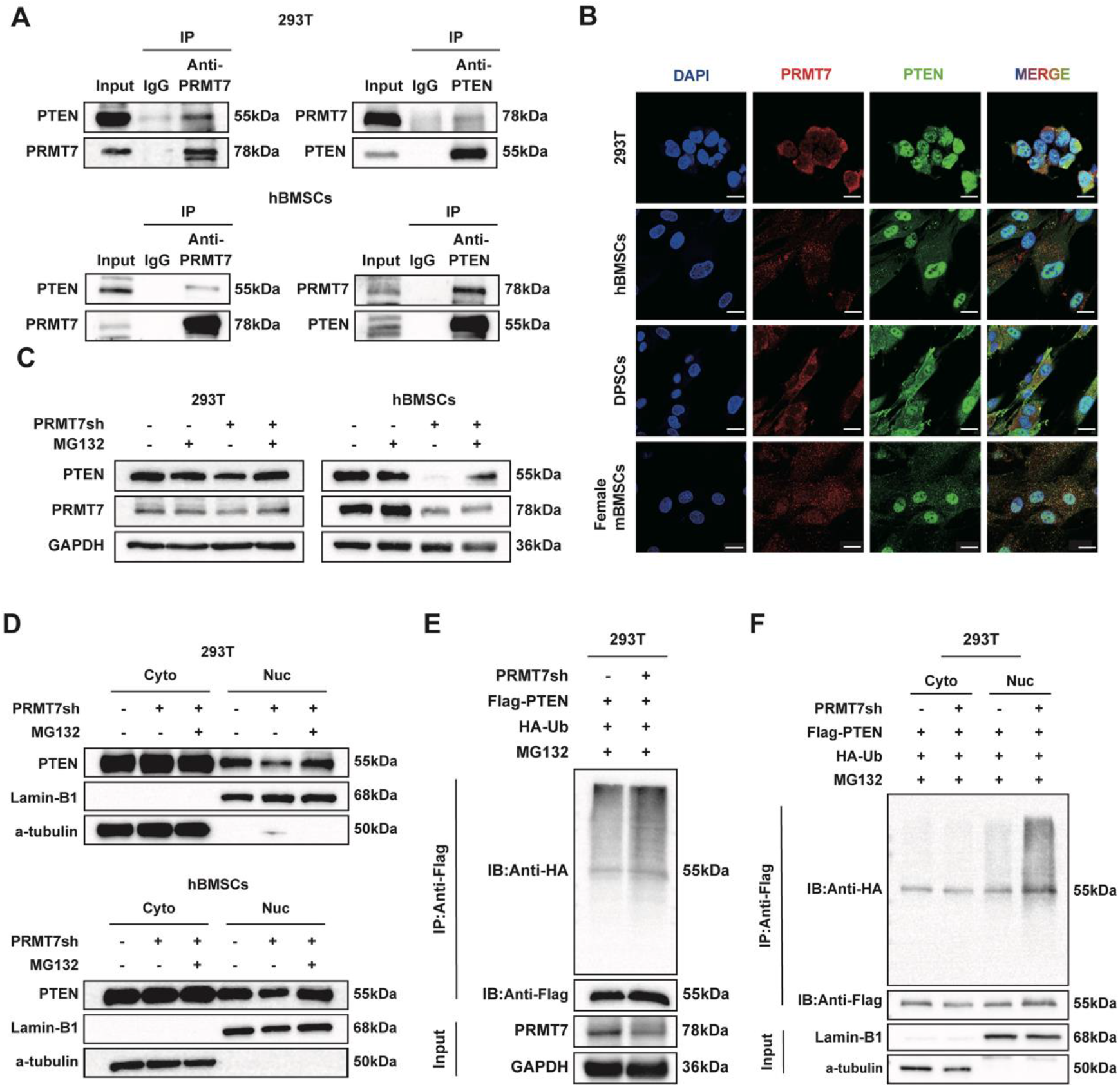
PRMT7 binds to and stabilizes PTEN in the nucleus. A. CoIP analysis showing the interaction of PRMT7 with PTEN in 293T and hBMSCs lysates with anti-PRMT7 and anti-PTEN antibody (pull-down with anti-PRMT7 or anti-PTEN antibody). IgG control blots confirming the specificity of the CoIP experiments. n = 3 biological replicates. B. Co-localization of PRMT7 and PTEN in 293T, hBMSCs, DPSCs and female mBMSCs. Scale bar: 20 μm. C. Representative Western blot images of PTEN in PRMT7sh and control cells. Cells were treated with or without 10 μM MG132 for 6 hours prior to harvesting. n = 3 biological replicates. D. Representative Western blot images of cytoplasmic and nuclear PTEN in PRMT7sh and control cells. Cells were treated with or without 10 μM MG132 for 6 hours prior to harvesting. n = 3 biological replicates. E. 293T cells were transfected with the indicated plasmid, followed by treatment with 10 μM MG132 for 6 hours. Subsequently, *in vivo* ubiquitination assays and western blot were conducted. F. 293T cells were transfected with the plasmids, followed by treatment with 10 μM MG132 for 6 hours. Subsequently, nuclear and cytoplasmic fractions were isolated for *in vivo* ubiquitination assays and analyzed by Western blot. n = 2 biological replicates.

Given that PRMT7 deficiency reduces both PTEN mRNA and protein levels, we aimed to determine whether the absence of PRMT7 also influences PTEN protein degradation, beyond its effect on transcription. Therefore, the proteasome inhibitor MG132 was added and the PTEN reduction induced by PRMT7 knockdown was reversed by MG132, indicating that PRMT7 affected PTEN stability through the ubiquitin-mediated protein system (Fig. 6C; Appendix Fig. 5B). A previous study showed that MG132 could only restore the protein level of nuclear PTEN, but not cytoplasmic PTEN. Therefore, we hypothesized that knockdown of PRMT7 could only reduce the expression of nuclear PTEN, but not cytoplasmic PTEN (*Ge et al., 2020*). This was confirmed by our nuclear-cytoplasmic separation assay, in which nuclear PTEN was significantly reduced while cytoplasmic PTEN remained unchanged after PRMT7 knockdown. Furthermore, only nuclear PTEN could be restored by MG132 (Fig. 6D; Appendix Fig. 5C). We also found that the effect of PRMT7 on the stability of nuclear PTEN via the ubiquitin-proteasome system is independent of PRMT7’s methyltransferase activity, indicating that this regulatory mechanism is methyltransferase-independent (Appendix Fig. 5D). Then, lentivirus, Flag-PTEN, and HA-Ub were added into 293T cells to examine the effect of PRMT7 on PTEN ubiquitination. The results showed that PTEN ubiquitination was increased after PRMT7 knockdown (Fig. 6E). Next, we purified 293T cell lysates for nuclear and cytoplasmic fractionation followed by IP with Flag agarose beads. We found that knockdown of PRMT7 only increased nuclear PTEN ubiquitination levels but not cytoplasmic PTEN (Fig. 6F). These results support that knockdown of PRMT7 mediates ubiquitylation of nuclear PTEN, which is responsible for its degradation.

### PTEN rescued bone loss in *Prmt7* CKO mice

To investigate the potential role of PTEN in PRMT7-regulated osteogenesis, we examined the function of PTEN in osteogenesis. Appendix Fig. 6A-B demonstrated increased PTEN protein and mRNA expression after osteogenic induction (Appendix Fig. 6A-B). ALP staining and quantification revealed a significant decrease in osteogenic differentiation ability following PTEN deletion with siRNA (Appendix Fig. 6C-E). Our findings demonstrate that PTEN facilitates the osteogenic differentiation of MSCs, aligning with previous studies that have shown PTEN’s critical role in the osteogenic differentiation of both BMMSCs and DPSCs (*Nowwarote et al., 2023; Shen et al., 2019*).

To verify whether PTEN could reverse the *Prmt7*-induced bone loss *in vivo*, we employed adeno-associated virus serotype 9 (AAV9), known for its bone targeting by systemic delivery (Yang et al., 2019), to carry PTEN into bone tissue. After 6 weeks of systemic injection in 6-week-old female *Prmt7*-CKO and control mice, the infection efficiency of AAV9-control/*Pten* was detected in major organs and bones of all experimental groups by optical imaging and Western blot of extracted BMSCs (Fig. 7A-B). H&E staining of liver sections showed that AAV9 had no significant effect on this major metabolic organ (Appendix Fig. 6F). Micro-CT analysis demonstrated that PTEN can reverse the bone loss caused by PRMT7 deletion. This phenotype was observed in the femur, skull, and mandible. Quantitative analysis of bone parameters indicated that BMD, BV/TV, and Tb. N were decreased in CKO mice and increased in AAV9-*Pten* groups, while Tb. Sp displayed an opposite trend (Fig. 7C-H). We also observed that pre-dentin was relatively wider in AAV-*Pten* groups compared to the control group, indicating that teeth biomineralization returned to normal levels (Fig. 7I). These results demonstrated that PTEN could reverse *Prmt7*-deficient induced bone loss in female mice.

**Figure 7.**
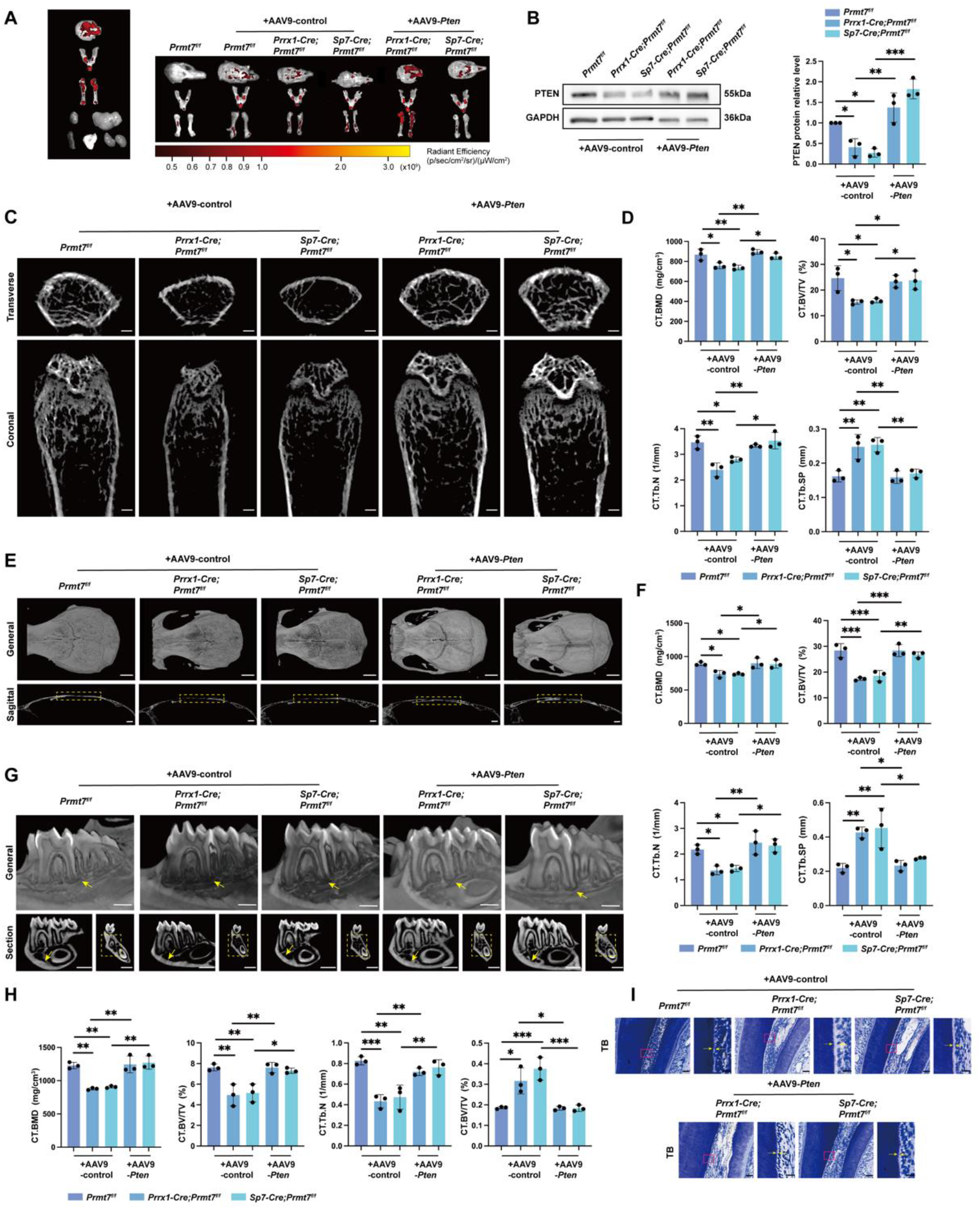
Bone loss in *Prmt7* CKO mice is mitigated by PTEN. A. *In vivo* fluorescence imaging was performed on the calvaria, mandible, femur, heart, liver, spleen, lung, kidneys (left), and bone tissues across all groups of mice six weeks post-lentivirus injection. Radiant efficiency is depicted using bar graphs, with measurements given in (p/sec/cm²/sr)/(µW/cm²). B. Western blot analysis (left) of PTEN in mBMSCs, isolated from female *Prmt7^f/f^*, *Prrx1-Cre; Prmt7^f/f^* and *Sp7-Cre; Prmt7^f/f^* mice six weeks following tail vein administration of AAV9-control or AAV9-*Pten*. Quantification (right) of relative PTEN levels normalized to GAPDH. C. Micro-CT analysis of femurs from 12-week-old female *Prmt7^f/f^*, *Prrx1-cre; Prmt7^f/f^*and *Sp7-cre; Prmt7^f/f^* mice after 6-week of AAV9-control or AAV9-*Pten* injection. The upper panel shows the cross-sectional view of the metaphysis; The lower panel shows the coronal view of the metaphysis. Scale bar: 500 μm. D. Quantitative analysis of bone parameters at the femoral metaphysis growth plate, including BMD, BV/TV, Tb. N, and Tb. Sp, obtained in C (n = 3 mice for each group). E. Micro-CT analysis of skull from 12-week-old female *Prmt7^f/f^*, *Prrx1-cre; Prmt7^f/f^*and *Sp7-cre; Prmt7^f/f^* mice after 6-week of AAV9-control or AAV9-*Pten* injection. The upper panel: 3D reconstruction of the whole skull; the lower panel: midsagittal view of the skull. The yellow dashed box indicates the calvarial region. Scale bars: 2 mm (upper) and 1 mm (lower). F. Quantitative analysis of bone parameters at the calvarial region, including BMD, BV/TV, Tb. N, and Tb. Sp, obtained in E (n = 3 mice for each group). G. Micro-CT analysis of mandible from 12-week-old female *Prmt7^f/f^*, *Prrx1-cre; Prmt7^f/f^*and *Sp7-cre; Prmt7^f/f^* mice after 6-week of AAV9-control or AAV9-*Pten* injection. The upper panel: 3D reconstructed sectional view of the mandible; the lower left panel: mesiodistal sectional view of the first molar; the lower right panel: coronal sectional view of the first molar. The yellow arrow at the upper panel points to the bone around the apical area of the distal root of the first molar; the yellow arrow at the lower left panel points to the bone wall at the lower border of the mandibular body; the yellow dashed box at the lower right panel indicates the bone between the furcation of the first molar and the mandibular canal. Scale bars: 1 mm. H. Bone parameters quantitative analysis of mandible, including BMD, BV/TV, Tb. N, and Tb. Sp, obtained in E (n = 3 mice for each group). I. Toluidine Blue staining of the dental tissues at the alveolar crest of the distal root of the first molar from 12-week-old female *Prmt7^f/f^*, *Prrx1-cre; Prmt7^f/f^* and *Sp7-cre; Prmt7^f/f^* mice after 6-week of AAV9-control or AAV9-*Pten* injection. The yellow arrow in the right points to the predentin, which is an enlarged view of the area in the pink box in the left. Scale bars: 50 μm (left) and 20 μm (right). All data are mean ± SD. (*, *P*<0.05, **, *P*<0.01, ***, *P<*0.001) (One-way ANOVA).

In addition, we performed an *in vitro* rescue experiment by extracting BMSCs from different strains of mice and adding *Pten*-overexpression or control plasmids. Subsequent assays validated PTEN as an intermediate target between PRMT7 and osteogenesis. ALP staining and quantification showed that *Pten* enhanced osteogenic differentiation, which was reduced by *Prmt7* deletion (Appendix Fig. 6G). This was further confirmed by the examination of protein and mRNA levels of RUNX2 (Appendix Fig. 6H-I). Taken together, these results imply that PTEN is essential for bone formation in *Prmt7*-deficient female mice.

## Discussion

Our study reveals that PRMT7 specifically enhances bone formation and regeneration in the axial and appendicular bones of female mice, with no impact on males, through the construction of the control and CKO mouse models. PRMT7 facilitates osteogenic differentiation of BMSCs by activating PTEN. Mechanically, PRMT7 increases H3R2me1 at the *PTEN* promoter and stabilizes nuclear PTEN in a methyltransferase-independent manner. Moreover, PRMT7 knockdown diminishes osteogenic differentiation, an effect reversible by PTEN re-expression.

Initially, PRMT7 was mistakenly identified as an enzyme that catalyzes the formation of symmetrical dimethylation products due to contamination with PRMT5 (*Feng et al., 2013; Bedford & Clarke, 2009*). For instance, a previous study reported that PRMT7 catalyzed H3R2me2s to enhance binding with WDR5 in euchromatin (*Migliori et al., 2012*). However, current studies have clearly demonstrated that PRMT7 can only catalyze MMA (*Structural Determinants for the Strict Monomethylation Activity by Trypanosoma brucei Protein Arginine Methyltransferase 7, 2014; Feng et al., 2014*), and the alteration of SDMA may be due to crosstalk with other PRMTs via secondary effects, which has not been fully confirmed (*Jain et al., 2017; Yang & Bedford, 2013*). Given PRMT7’s ability to produce only MMA, we focused on its direct effect on H3R2. PRMT7 has been reported to be responsible for H3R2me1 at the promoter of *Rap1a*, affecting monocyte characteristics (*Günes Günsel et al., 2022*). H3R2me1 is associated with gene transcriptional activation, similar to H3R2me2s but opposite to H3R2me2a (*Kirmizis et al., 2007; Guccione et al., 2007; Hyllus et al., 2007*). We confirmed that H3R2me1 serves as a substrate of PRMT7 in BMSCs, regulating osteogenic differentiation. The loss of PRMT7 decreased the global expression level of H3R2me1 in MSCs and reduced its presence at the promoter of the target gene *PTEN*, thereby impacting its transcription and osteogenesis. Notably, a PRMT7 enzymatic dead mutant could not rescue the reduction in global H3R2me1 levels and its presence at the *PTEN* promoter caused by PRMT7 knockdown. Consequently, PRMT7 can be targeted in cell-mediated regenerative medicine to regulate osteogenic differentiation, influencing bone formation and regeneration.

The regulation of PTEN is governed by a multitude of molecular mechanisms. In addition to genetic alterations, PTEN is modulated by epigenetic modifications, microRNA regulation, post-translational modifications, and interactions with other proteins, forming an intricate regulatory network that ensures its proper physiological function (*Song et al., 2012*). Several studies have elucidated the regulatory influence of the PRMT family on PTEN via distinct mechanisms. PRMT5 represses PTEN expression at the transcriptional level in Glioblastoma Neurospheres (GBMNS) by binding to its promoter (*Banasavadi-Siddegowda et al., 2017*). PTEN protein undergoes ADMA at R159 through its interaction with PRMT6, influencing mRNA alternative splicing (*Feng et al., 2019*). A recent finding indicated that PRMT7 associates with PTEN, inducing monomethylation, thereby inhibiting gastric cancer (GC) cell proliferation and migration (*Wang et al., 2023*). Mechanistically, our research introduces novel insights into the PRMT-PTEN axis. PRMT7 enhances H3R2me1 levels at the PTEN promoter, activating PTEN expression transcriptionally. Additionally, PRMT7 modulates PTEN ubiquitination through a non-methyltransferase-dependent pathway, contributing to PTEN degradation via the ubiquitin-proteasome system. Phenotypically, our study highlights the critical role of the PRMT7-PTEN interaction in bone tissue, extending beyond its oncological implications.

Unlike cytoplasmic PTEN, which primarily functions in a phosphatase-dependent manner, nuclear PTEN performs various roles independent of its phosphatase activity (*Lindsay et al., 2006*). For example, the loss of nuclear PTEN can lead to more aggressive tumors by regulating genomic stability and the cell cycle (*Shen et al., 2007; Hou et al., 2019; Puc et al., 2005*). Prior research has demonstrated that the nuclear export of PTEN, rather than the inhibition of its phosphatase activity, compromises the activation of the Anaphase Promoting Complex/Cyclosome (APC)-CDC20like protein 1 (CDH1) complex, consequently reducing its tumor-suppressive function (*Song et al., 2011*). Considering the distinct roles of nuclear versus cytoplasmic PTEN and the pivotal significance of nuclear PTEN, it is imperative to investigate the regulatory mechanisms and functional pathways governing these differences. Emerging research has highlighted that the knockdown of NEDD4-1 and WWP2, two oncogenic ubiquitin ligases, selectively augments cytoplasmic PTEN levels without affecting nuclear PTEN (*Wang et al., 2007; Maddika et al., 2011*). Conversely, the E3 ligase FBXO22 specifically ubiquitinates nuclear PTEN, thereby impacting its stability (*Ge et al., 2020*). Furthermore, there is ongoing debate regarding the relative stability of nuclear versus cytoplasmic PTEN, potentially due to differences in endogenous versus exogenous sources of PTEN protein. Overexpression of exogenous PTEN can lead to protein instability and challenges in nuclear localization (*Ge et al., 2020; Trotman et al., 2007b*).

Our research demonstrates that PRMT7, beyond its role as a methyltransferase, acts as a novel stabilizer of nuclear PTEN, enhancing its stability without affecting cytoplasmic PTEN and thereby promoting osteogenesis. Given that PRMT7 lacks E3 ubiquitin ligase activity, we propose that PRMT7 might function as an adaptor protein, facilitating the recruitment of an E3 ubiquitin ligase to PTEN to mediate its ubiquitination. This hypothesis necessitates further exploration. In a word, our results provide new insights into the relative instability of nuclear PTEN and expand the functional role of nuclear PTEN in stem cell biology.

One intriguing aspect of our study is that PRMT7 promotes osteogenesis exclusively in female mice, without affecting male mice. Upon PRMT7 knockdown, PTEN expression levels decreased solely in female mice, with no significant change in males. This suggests PTEN may be a crucial mediator of the sex-specific bone phenotype driven by PRMT7. Numerous studies highlight the pervasive nature of sex differences across species, tissues, and biological processes (*Yang et al., 2006; Melé et al., 2015; Gershoni & Pietrokovski, 2017; Wells, 2007*). These differences may be gene activity-related and induced. While gene expression on sex chromosomes is a major source, the sex-biased expression of autosomal genes, often tissue-specific, is also noteworthy (*Naqvi et al., 2019*). PTEN, located on chromosome 10 in humans and chromosome 19 in mice, exhibits sex-dependent tissue specificity, encompassing adipose, musculature, central nervous system, thyroid gland, and hepatic tissue (*Zhang et al., 2005; Samaan et al., 2015; de Mello et al., 2020; Metcalfe & Stewart, 2023; Antico-Arciuch et al., 2010; Anezaki et al., 2009*). For example, a high-fat diet raises AEBP1 levels in white adipose tissue and reduces PTEN expression in female mice, while male mice show no significant changes in AEBP1 or PTEN (*Zhang et al., 2005*). These differences are primarily linked to sex hormone levels, with several studies indicating a close correlation between hormone/hormone receptor expression and PTEN levels (*Chen et al., 2018; Li et al., 2023b; Vilgelm et al., 2006*). In endometrial cancer, estrogen binds to ER, increasing miR-200c levels and inhibiting PTEN and PTENP1, which activates the PI3K-AKT pathway (*Chen et al., 2018*). Although our RNA-seq results showed no significant differences in estrogen and receptor levels between female CKO and control mice, the higher estrogen levels in females compared to males may explain the PTEN alterations observed exclusively in female mice, warranting further investigation (*Almeida et al., 2017*). Taken together, our study highlights gender-specific differences in PTEN expression in bone tissue, reinforcing evidence of its modulation by sex.

In conclusion, these findings reveal that PRMT7 influences osteogenesis via both methyltransferase-dependent and -independent pathways, highlighting its essential physiological role in bone formation and establishing it as a key target for bone repair and regenerative therapies.

## Methods

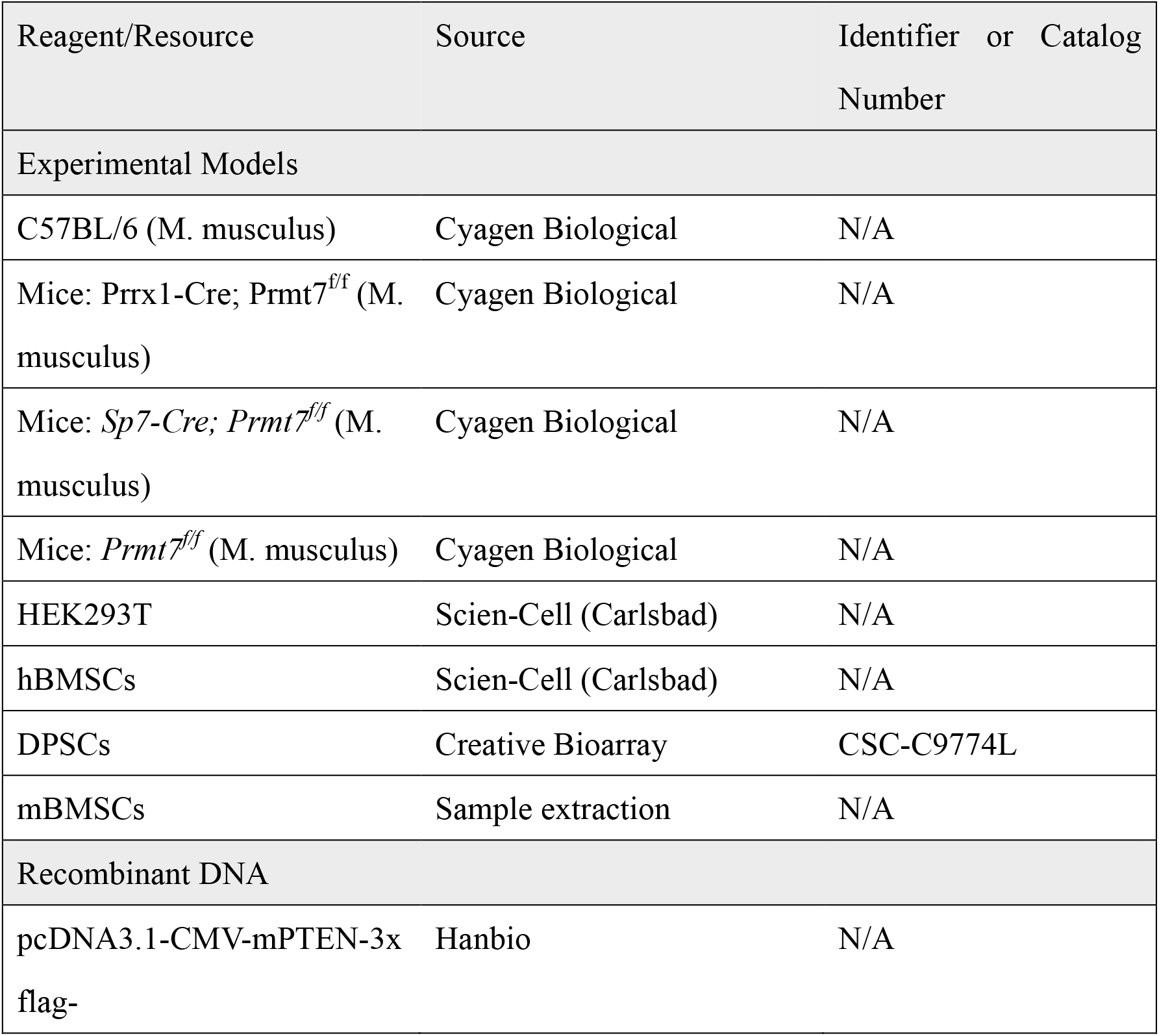
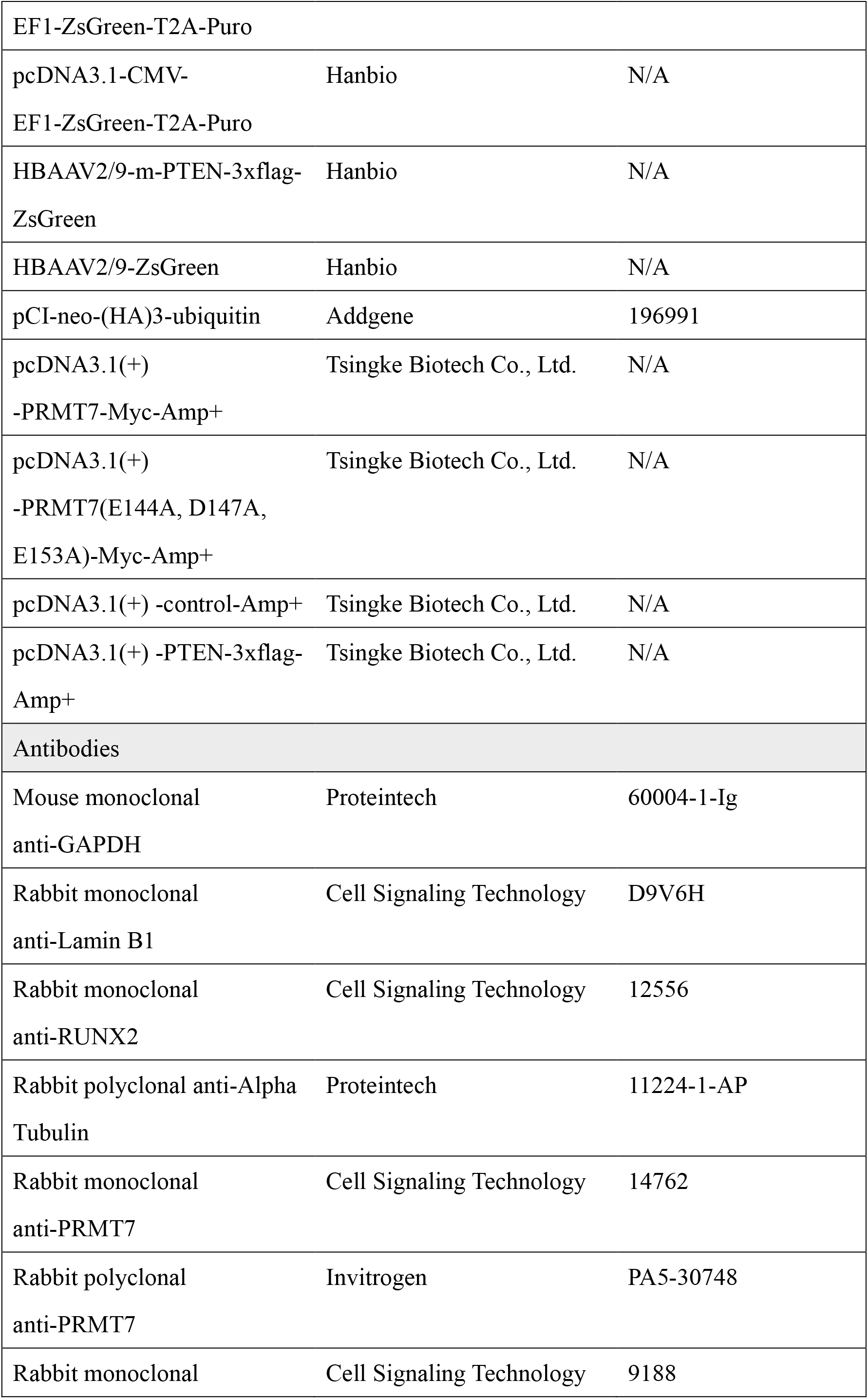
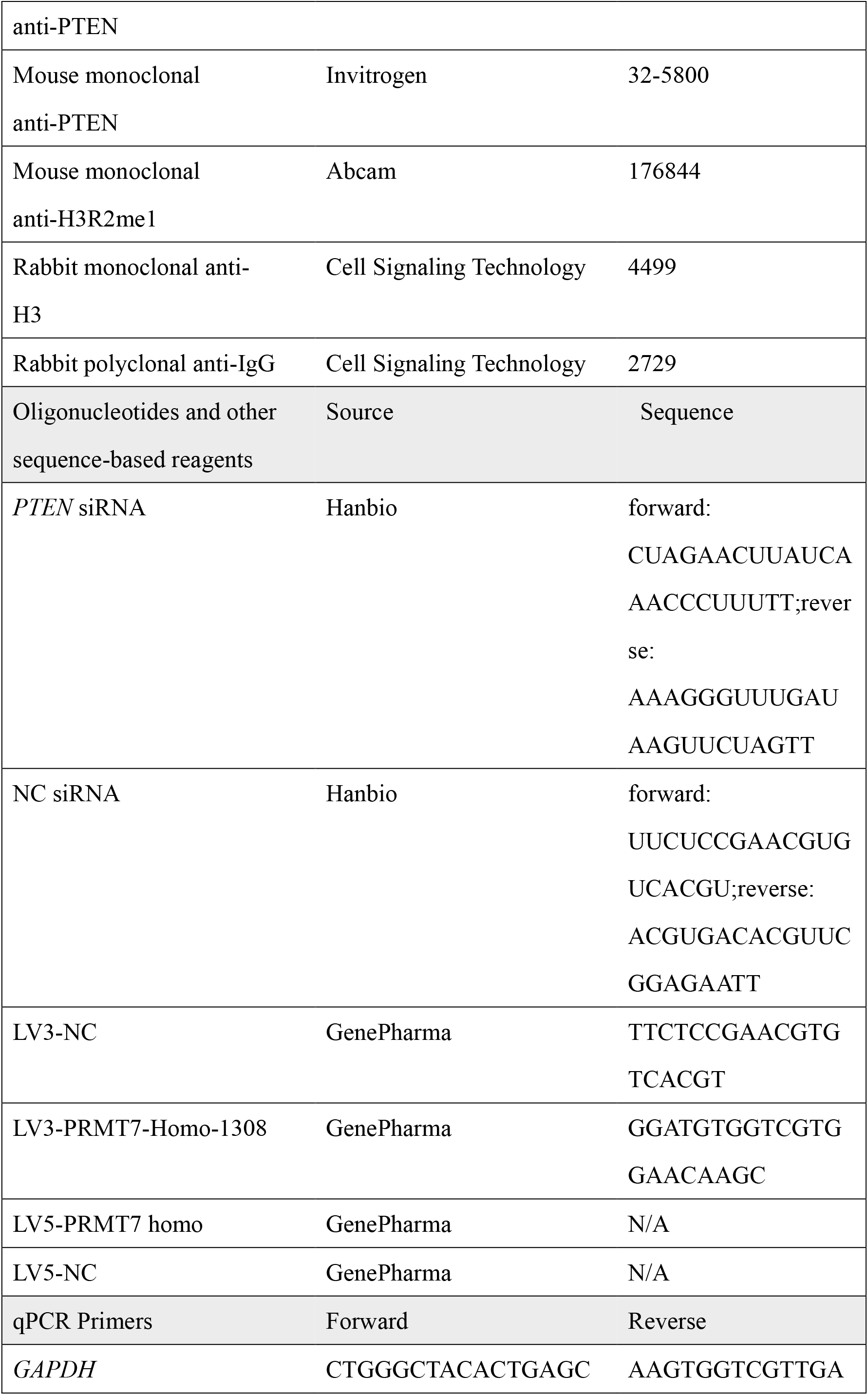
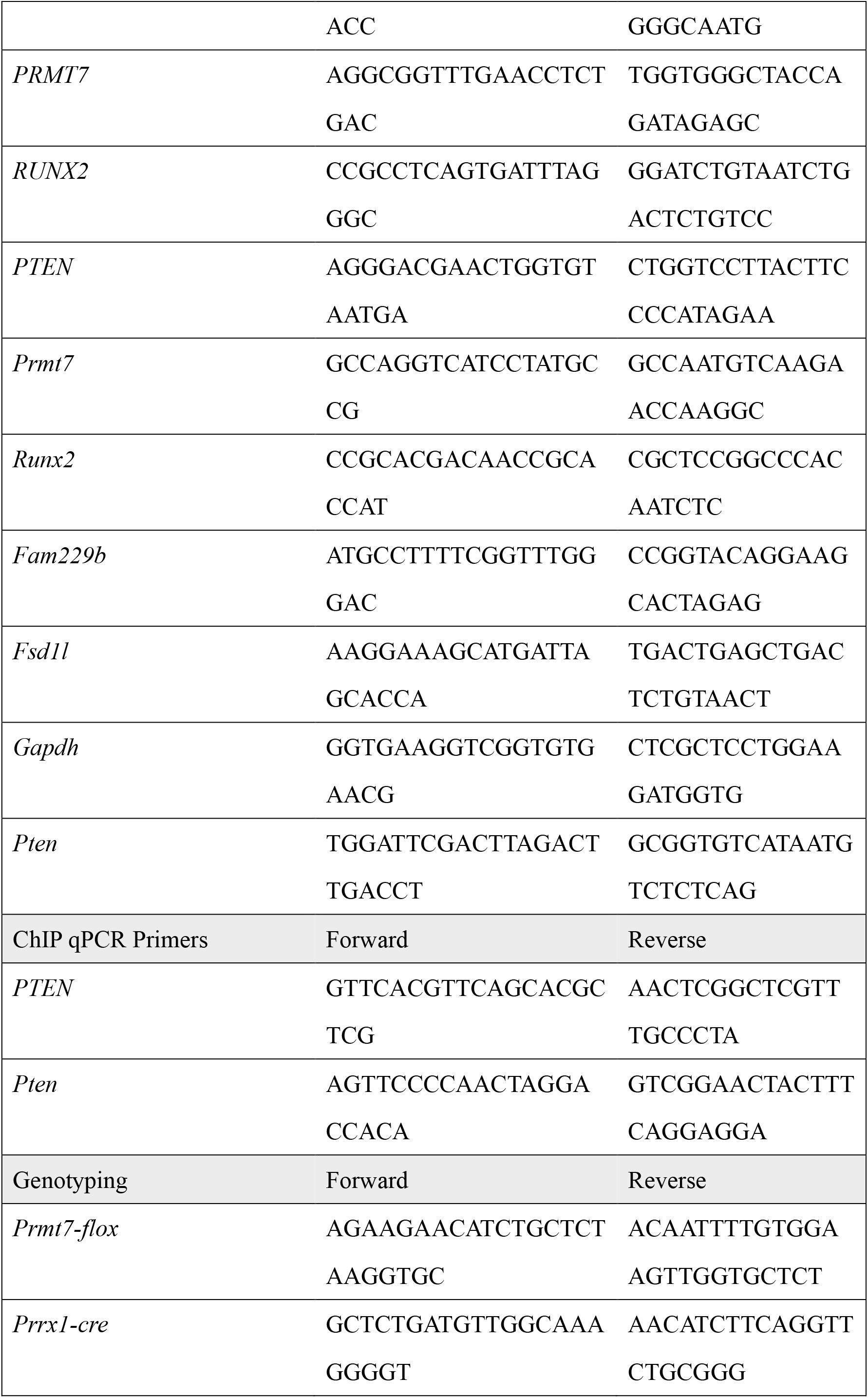
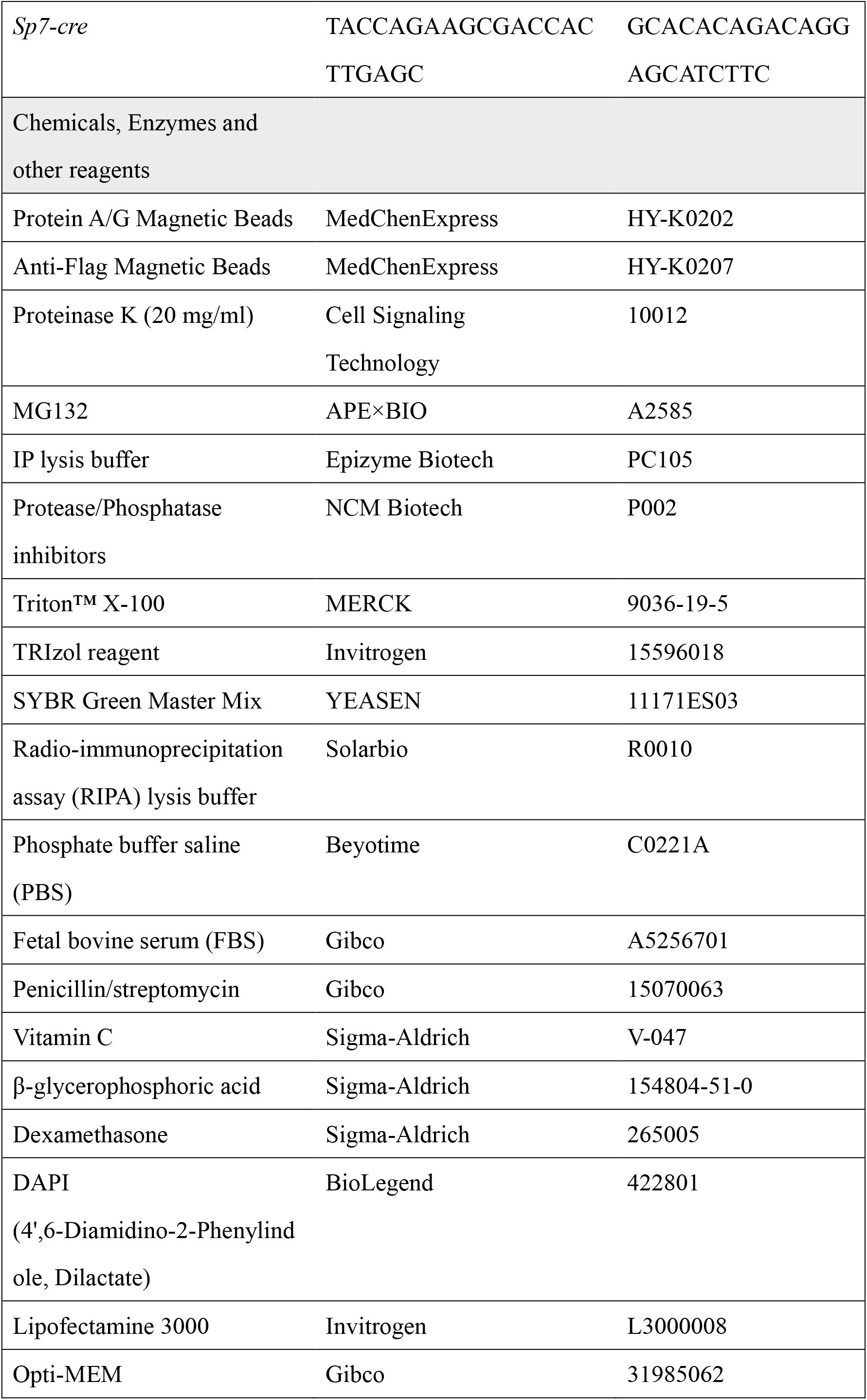
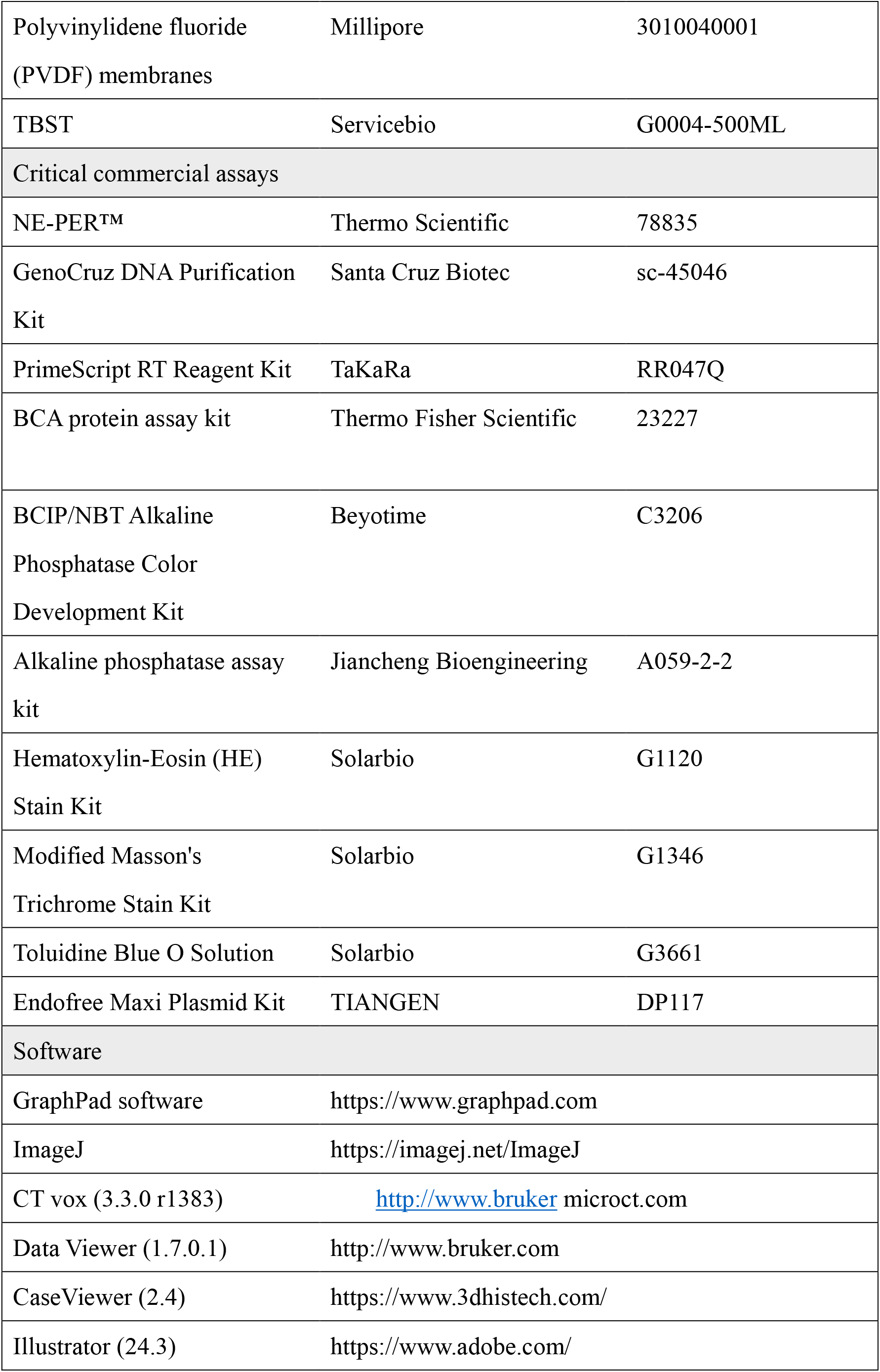
Reagents and Tools Table

### Experimental animal

All mice used were C57BL/6J genetic background and were grown in a specific pathogen free (SPF) environment with a 12-hour day-night cycle. *Prmt7^f/f^*mice were obtained from Biocytogen Co., Ltd. (Beijing, China) by adding LoxP sites on either side of exons 3 and 4 of *Prmt7*. These *Prmt7^f/f^* mice were then mated with *Prrx1-cre* and *Sp7-cre* mice (also from Biocytogen Co., Ltd.) individually to obtain *Prrx1-cre; Prmt7^f/f^* and *Sp7-cre; Prmt7^f/f^*mice. Genotype identification was performed, and the related primers are listed in Appendix Table 1. For colocalization immunofluorescence experiments, we used 6-week-old female mice. For *in vivo* imaging experiments, 8-week-old female mice were used as controls. All animal experimental procedures were approved by the Institutional Animal Care and Use Committee of Peking University Health Science Center.

### Defect model

For the tibial defect model, a 1mm diameter round bur was used to create a defect in the right proximal tibia of the mouse. The wound was washed and sutured, and samples were collected 10 days later. For the skull defect model, an annular drill with an inner diameter of 2mm was used to create a defect in the middle of the calvarium of the mouse. The wound was debrided and closed. For the mandible defect model, the skin was cut along the midline between the angle of the mouth and the tragus of the ear. The masseter muscle was exposed, and the blood vessels, nerves, and muscles were separated to locate the mandibular bone wall. A 1.5mm ball drill was then used to create the defect, and the mice were sacrificed after 2 months.

### Micro-CT and quantitative analysis

The collected bone tissue samples were fixed in 4% paraformaldehyde for 2 days and subsequently scanned using a SkyScan 1276 (Bruker, China) with a voltage of 80 kV, a current of 200 μA, a spatial resolution of 6 μm, and a pixel value of 2048 x 2048. We then used the NRECON software to reconstruct the images. The DATAVIEWER, CTVOX, and CTAN software can be used to obtain 2D images, 3D images, and quantitative analysis parameters of the bone samples, respectively.

## Whole-body skeleton staining

Collected samples were removed with forceps for skin and viscera, then fixed in 95% ethanol for 24 hours. The samples were then transferred to Alcian Blue solution for cartilage staining for three days. Decolorization was performed with 95% ethanol for three days. The specimens were then immersed in 2% KOH for 24 hours to achieve transparency, followed by a 24-hour counterstain in 0.015% Alizarin Red solution (Sigma A3757). Finally, the samples were placed in a mixture of 20% glycerin and 1% KOH for preservation.

### Histological staining

Bone samples and liver were fixed with 10% paraformaldehyde for 48 hours and decalcified with 10% EDTA-2Na solution (pH 7.4, Baoya) for 2 weeks (decalcification was not required for the liver). The samples were subsequently dehydrated overnight using a gradient sucrose solution. Frozen sections were then prepared using a freezing microtome (Leica CM1950) after OCT embedding. The slices were stained according to the different kits from Solarbio (HE Stain Kit, Masson’s Stain Kit, and Toluidine Blue O Solution).

### Cell extraction, culture, and *in vitro* assay

Mouse BMSCs were flushed from the bone marrow of 6-week-old CKO and control mice and then seeded in 6cm or 10cm dishes. All cells were grown in a proliferation medium (PM) containing α-minimum essential medium (α-MEM) (Gibco), 10% fetal bovine serum (FBS) (Gibco), and 1% v/v penicillin/streptomycin (P/S) (Gibco) at 37°C and 5% CO2. The medium was changed every two days.

For different experimental purposes, we performed osteogenic induction experiments when the cells reached 90% confluency. Osteogenic induction was conducted using an osteogenic induction medium that included α-MEM, FBS, P/S, 10 nM dexamethasone (Sigma), 200 μM vitamin C (Sigma), and 10 mM β-glycerophosphate (β-GP, Sigma). The induction solution was changed every two days. ALP staining (Beyotime, Shanghai, China) and quantification (Nanjing Jiancheng Bioengineering Institute, Nanjing, China) were performed after 7 days of osteogenic induction according to the instruction manual.

### Lentiviral transduction and plasmid, siRNA transfection

Cells were transduced with PRMT7sh, PRMT7, and negative control lentivirus (Gene Pharma, Jiangsu, China) for 24 hours. Then, the medium was replaced, and puromycin (1 mg/ml, Sigma-Aldrich) was added. After 48 hours, transduction efficiency was assessed using fluorescence imaging, Western blot, and qRT-PCR. Plasmid DNA and siRNA were transfected into cells using Lipofectamine 3000 (Invitrogen). Cells were seeded 24 hours before transfection. DNA/siRNA-lipid complexes were formed in Opti-MEM(Gibco) and added to the cells. Cells were harvested 48 hours post-transfection, and transfection efficiency was assessed using western blot and qRT-PCR.

### Total Protein extraction and western Blot

Cells were collected and placed on ice, then homogenized in RIPA buffer (PC102, Epizyme Biotech, Shanghai, China) with protease and phosphatase inhibitors (NCM Biotech, Jiangsu, China). The homogenates were incubated on ice for 30 minutes and centrifuged at 14,000 rpm for 15 minutes at 4°C. Supernatants were collected, and protein concentrations were measured using a BCA protein assay kit (Thermo Fisher Scientific, Waltham, MA, USA). Equal amounts of protein (20 μg) were separated by SDS-PAGE and transferred to a PVDF membrane (Millipore, Burlington, MA, USA). The membrane was blocked with 5% skim milk in TBST (Servicebio, Hubei, China) for 1 hour at room temperature, then incubated with primary antibodies overnight at 4°C. After washing with TBST, the membrane was incubated with HRP-conjugated secondary antibodies for 1 hour at room temperature. Bands were visualized using enhanced chemiluminescence (ECL, Epizyme Biotech, Shanghai, China) and an imaging system. Protein intensities were quantified using ImageJ.

### Nuclear and cytoplasmic protein extraction

Nuclear and cytoplasmic proteins were extracted using the Thermo Scientific Nuclear and Cytoplasmic Extraction Reagents (NE-PER™) kit according to the manufacturer’s instructions. Briefly, cells were harvested and resuspended in Cytoplasmic Extraction Reagent I (CER I), incubated on ice, and then treated with Cytoplasmic Extraction Reagent II (CER II). The cytoplasmic fraction was collected after centrifugation. The remaining pellet was resuspended in Nuclear Extraction Reagent (NER) and incubated on ice with intermittent vortex. The nuclear fraction was collected following centrifugation. Both fractions were stored at -80°C until further analysis.

### RNA Extraction and qPT-PCR

Total RNA was extracted from samples using TRIzol reagent (Invitrogen) following manufacturer’s instructions. RNA concentration and purity were measured with a NanoDrop spectrophotometer (Thermo Fisher Scientific). cDNA was synthesized from 1 µg of total RNA using a cDNA synthesis kit (Accurate, Hunan, China). PCR was carried out using SYBR Green PCR Master Mix (Vazyme Biotech, Nanjing, China) on a real-time PCR system. The PCR conditions were 95°C for 3 minutes, followed by 40 cycles of 95°C for 10 seconds and 60°C for 30 seconds. Gene expression levels were normalized to *GAPDH*.

### RNA sequencing

Total RNA was extracted from female *Prmt7^f/f^*, *Prrx1-Cre; Prmt7^f/f^*, and *Sp7-Cre; Prmt7^f/f^* mBMSCs using TRIzol reagent (Invitrogen). RNA quality was assessed with a NanoDrop spectrophotometer and an Agilent 2100 Bioanalyzer, with RIN > 7. mRNA was isolated, fragmented, and reverse transcribed into cDNA. After end repair and adapter ligation, cDNA libraries were PCR-enriched and sequenced on an Illumina platform.

### ChIP-qPCR

Cells were cross-linked with 1% formaldehyde and quenched with 0.125 M glycine. Chromatin was sheared into 200-500 bp fragments by sonication. Immunoprecipitation was performed using antibodies against the target protein and protein A/G magnetic beads (MCE). After washing, cross-links were reversed at 65°C overnight and treated with proteinase K. DNA was purified and analyzed by qPCR with specific primers. Data were normalized to input DNA and analyzed using the ΔΔCt method.

### Immunoprecipitation (IP)

Cells were lysed in IP lysis buffer (Epizyme Biotech, Shanghai, China) containing protease inhibitors. Protein lysates were pre-cleared with protein A/G magnetic beads, followed by incubation with antibodies overnight at 4°C. Protein complexes were captured using protein A/G or anti-Flag magnetic beads, washed, and eluted in the SDS sample buffer. Eluates were subjected to SDS-PAGE and analyzed by Western blotting using relevant antibodies.

### *In vivo* ubiquitination assay

To assess the *in vivo* ubiquitination of PTEN, HEK293 cells were transfected with plasmids encoding HA-tagged ubiquitin, Myc-tagged PRMT7, and FLAG-tagged PTEN. Forty-eight hours post-transfection, cells were treated with 10 μM MG132 (MCE) for 6 hours to inhibit proteasomal degradation. Cells were then harvested and lysed in IP lysis buffer or Nuclear and Cytoplasmic Extraction Reagents (NE-PER™) kit supplemented with protease and phosphatase inhibitors. The lysates were clarified by centrifugation at 14,000 rpm for 15 minutes at 4°C. The supernatants were subjected to immunoprecipitation using Anti-Flag Magnetic Beads overnight at 4°C. Immunoprecipitated proteins were washed extensively with RIPA buffer and eluted by boiling in SDS-PAGE sample buffer. Western blot analysis was performed using anti-HA and anti-FLAG antibodies to detect ubiquitinated PTEN.

### Co-localization assay

Cells were fixed in 4% paraformaldehyde, permeabilized, and blocked. Double immunofluorescence staining was performed using antibodies against PRMT7 and PTEN, followed by appropriate secondary antibodies conjugated with fluorophores. Images were acquired using a fluorescence microscope (Olympus FV3000).

### Intravenous injection and *in vivo* Imaging

*Prmt7^f/f^*, *Prrx1-Cre; Prmt7^f/f^* and *Sp7-Cre; Prmt7^f/f^* female mice aged 6 weeks were prepared for intravenous injection via the tail vein. Before injection, the mice were gently restrained to expose the tail vein. A 27-gauge needle attached to a syringe containing AAV-*Pten* or AAV-control was inserted into the vein, ensuring minimal discomfort and proper injection technique to avoid leakage or vascular damage. After injection, the mice were monitored for any immediate adverse reactions.

For *in vivo* imaging, mice were anesthetized via inhalation of isoflurane to minimize motion artifacts. Imaging was performed using bioluminescence imaging. Imaging settings were optimized for sensitivity and resolution, and data acquisition times were standardized across all experiments to ensure consistency. Post-imaging, mice were allowed to recover under controlled conditions with access to food and water. This protocol adheres to institutional guidelines for the humane use of animals in research and was approved by the Institutional Animal Care and Use Committee at Peking University.

### Statistical Analysis

All statistical analyses were performed using GraphPad Prism version 9.0 (GraphPad Software, San Diego, CA, USA). Data were expressed as mean ± standard deviation (SD) for continuous variables and as frequencies and percentages for categorical variables. Normality of data distribution was assessed using the Shapiro-Wilk test. For normally distributed data, comparisons between groups were made using the independent samples t-test or one-way ANOVA followed by post hoc Tukey’s test. For non-normally distributed data, the Mann-Whitney U test or Kruskal-Wallis test followed by Dunn’s post hoc test was employed. A p-value of less than 0.05 was considered statistically significant.

## Acknowledgements

We thank Luzheng Xu and Qilong Wang from the Peking University Medical and Health Analysis Center for their assistance with Micro CT images scanning and processing.

